# A combined biochemical and cellular approach reveals Zn^2+^-dependent hetero- and homodimeric CD4 and Lck assemblies in T cells

**DOI:** 10.1101/2022.12.17.520849

**Authors:** Anna Kocyła, Aleksander Czogalla, Inga Wessels, Lothar Rink, Artur Krężel

## Abstract

The CD4 or CD8 co-receptors’ interaction with the protein-tyrosine kinase Lck is widely accepted as the initiator of the tyrosine phosphorylation cascade leading to T-cell activation. These co-receptors potentially enhance T-cell antigen sensitivity, but how they function is still debated. A critical question is: to what extent are co-receptors and signal-initiating Lck coupled? Our contribution concerns the small – but indispensable for CD4- and CD8-Lck formation – element Zn^2+^. The intracellular Zn^2+^ pool is strictly buffered but undergoes dynamic changes, also reported during T-cell activation. Furthermore, the identical Zn^2+^-binding cysteinyl residues may alter co-receptor dimerization or heterodimerization with Lck. Following initial research demonstrating a significant difference in the affinity of Zn^2+^ to CD4 and CD4-Lck in solution, we combined biochemical and cellular approaches to show that fluctuations of buffered Zn^2+^ in physiological ranges indeed influence Zn(CD4)_2_ and Zn(CD4)(Lck). This conclusion was supported by the simulation of complexes’ equilibria, demonstrating that Zn^2+^ changes can alter the molar ratio between those complexes. In T cells, increased intracellular free Zn^2+^ concentration causes higher CD4 partitioning in the plasma membrane by a still unknown mechanism. We additionally found that CD4 palmitoylation decreases the specificity of CD4-Lck formation in the reconstituted membrane model, suggesting that this reversible modification may also be involved. Our findings help elucidate co-receptor-Lck coupling stoichiometry and demonstrate that intracellular free Zn^2+^ has a major role in the interplay between CD4 dimers and CD4-Lck assembly.

## Introduction

Activation of the αβ T-cell receptor (TCR) is mediated by the CD3 complex and supported by CD4 or CD8 co-receptors that extracellularly bind to MHC-II or MHC-I, respectively, and form the ternary TCR-MHC-CD4/CD8 complex (Fig. 1A) (1, 2). Inside the T cell, signal transduction is initiated by Src-family kinases, such as Lck (lymphocyte-specific protein-tyrosine kinase), which phosphorylates CD3 associated with a TCR complex. While co-receptors strongly enhance T-cell responses that are potent amplifiers of signaling events, their mechanistic mode of action is still a matter of debate (1-5). Lck interacts with the cytoplasmic tail of the co-receptor, thereby approximating the ligand-bound TCR-CD3 complex to initiate the phosphorylation cascade (6). However, it has been shown that a pool of Lck freely diffusing within the membrane is also involved (4, 7, 8). The two-stage co-receptor recruitment model suggests that the function of a co-receptor is modulated by TCR complex triggering, but not the other way around. The TCR complex is initially and partially phosphorylated by free Lck, followed by the recruitment of the co-receptor-Lck complex to the TCR-MHC complex (4, 7-9). Interestingly, even though CD4 and CD8 co-receptors deliver Lck to the TCR complex (10, 11), the effect of CD4 on the affinity and half-life of the TCR-MHC II complex was shown to be negligible (12-14). CD4-MHC II interaction is orders of magnitude weaker than typical T-cell/APC (antigen presenting cell) peptide interactions, suggesting that MHC has a very poor ability to recruit co-receptors (14).

**Figure 1.**
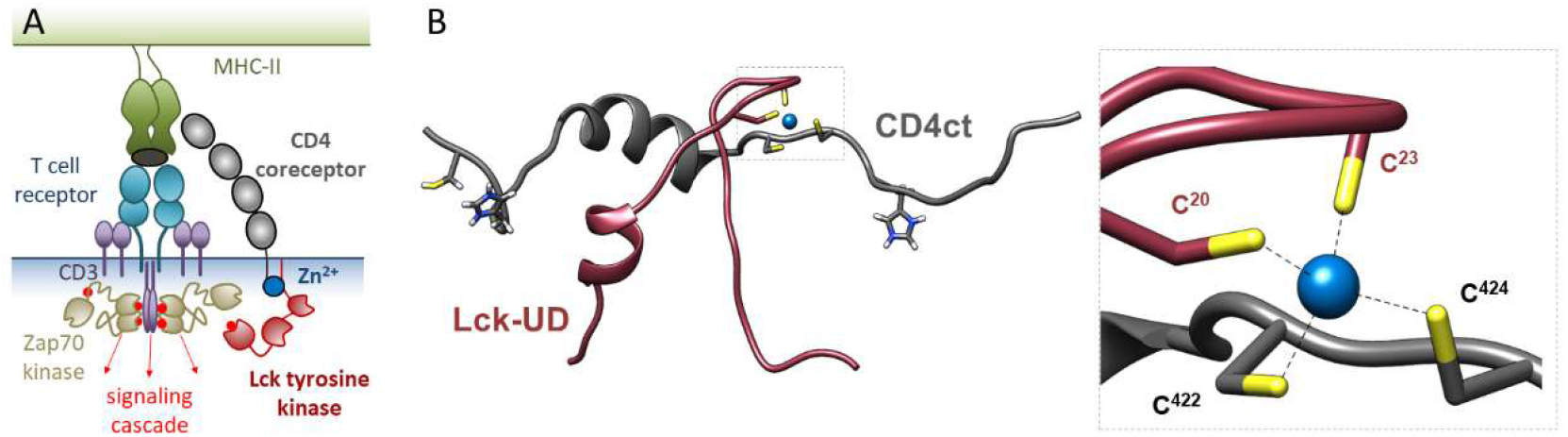
Architecture of zinc clasp assembly. A) Schematic representation of CD4-Zn^2+^-Lck complex formed in the T cell. Green and blue areas represent an antigen presenting cell and a T cell, respectively. B) NMR structure of zinc clasp (PDB: 1Q68) (19). Dashed box represents interprotein Zn^2+^ coordination environment in zinc clasp. Wine and gray cysteinyl residues come from CD4 and Lck molecules, respectively.

Accumulating data about independent signal transduction by CD4/CD8-Lck complexes resulted in the identification of a 32-amino-acid-long N-terminal domain of Lck, which is necessary and sufficient to interact with the 38-amino-acid-long C-terminal tail of CD4 (Fig. 1A). Within the domains four cysteine residues were found critical, together with a zinc ion (Zn^2+^) that links the CD4 and Lck units (15-18). Structural investigations of Zn(CD4/CD8)(Lck) complexes showed that Zn^2+^ is bound with micromolar affinity in a tetrahedral geometry within a compact fold called zinc clasp assembly (Fig. 1B) (19). However, further competition studies demonstrated much higher, subnanomolar Zn^2+^ affinity to CD4 that matches well free Zn^2+^ concentration (([Zn^2+^]_free_) fluctuations occurring in the cell, suggesting that zinc clasp assembly might be regulated in a Zn^2+^-dependent manner (20-22). Indeed, a significant rise in the intracellular free Zn^2+^ a minute after TCR triggering was observed (23). In the cell, Zn^2+^ ions are mostly released from intracellular stores, and it is known to modulate kinase/phosphatase balance in triggered signaling pathways (24-31). The potentiating effect of Zn^2+^ on T-cell activation was corroborated in T cells from human subjects supplemented with Zn^2+^ (24). However, the effects of changes in intracellular zinc homeostasis may be either stimulatory or inhibitory (31, 32).

T-cell activation is supported by membrane lipids that form specialized domains recruiting or excluding signaling proteins, e.g., immune synapse and protein-based clusters (33-37). The main protein modification guiding proteins to the membrane subdomain(s) is the reversible palmitoylation that occurs at the membrane-proximal cysteinyl residues. After T-cell stimulation, newly palmitoylated proteins emerge, thus lately, the palmitome started to be considered as a regulatory mechanism in T-cell signaling (38, 39). CD4 and Lck were identified in distinct membrane domains and their localization was strictly related to their palmitoylation (40-42). However, the local concentrations of CD4, Lck, and free Zn^2+^ additionally regulate the quantity of CD4 and Lck complexes (21). It means that not only free Zn^2+^ fluctuations but a local change of CD4 and Lck concentrations affects functional zinc clasp assembly formation. This is associated with palmitoylation, but also with the expression efficiency of co-receptors and Lck that differ, and their changes may affect the zinc clasp assembly (43-47).

Although the roles of both CD4/CD8 and Lck in thymic development and T-cell activation have been thoroughly investigated, the role of CD4/CD8-Lck complexes and mechanisms of their action within the immune responses are still debated (5). The latest reports suggest that in addition to initiating or augmenting the TCR response, they can also affect antigen affinity, which makes their mode of action more puzzling (48). Considering the importance of the CD4 and Lck interaction in the immune response, here we complement it with a Zn^2+^-dependent mode of their action. Three different approaches were undertaken: (i) in-solution studies using model peptides, (ii) membrane reconstitution of zinc clasp assembly, and (iii) T-cell studies with overexpression of CD4 and Lck where intracellular free Zn^2+^ availability was controlled. Experiments using artificial membranes and a model cell line showed that cellular Zn^2+^ fluctuations can induce zinc clasp assembly, thus making intracellular free Zn^2+^ concentrations a limiting factor. Moreover, our study demonstrated that the palmitoylation of CD4 affects Zn(CD4)(Lck) complex formation and confirmed the physiological relevance of Zn^2+^ in the plasma membrane status of CD4. Our data open new directions to the understanding of mechanistic details of TCR activation.

## Results

### Reconstitution of Zn^2+^-dependent interaction of CD4 and Lck

Previous reports showed that the formation of the zinc clasp assembly occurs at a subnanomolar range of buffered free Zn^2+^ concentration indicating high stability of the heterodimer (Zn(CD4)(lck)). A similar approach showed that homodimerization of the CD4 cytoplasmic tail occurs at a higher free Zn^2+^ concentration, showing that such a complex is significantly less stable (21, 49). To examine whether the zinc clasp assembly may occur in a more cellular environment as well, the interaction was reconstituted using artificial membranes at [Zn^2+^]_free_ controlled conditions provided by zinc competitors buffering its free concentration from 10^−14^ to 10^−9^ M (Figure 2A, SI). The C-terminal cytoplasmic domain of CD4 (CD4ct) and the N-terminal unique domain of Lck (Lck-UD) were myristylated to provide their docking (Fig. 2A, 2B). Fluorescent modifications at their C-terminus were introduced to track the assembly via FRET phenomena (Fig. 2B, 3A). These modifications enabled myrCD4ct(FAM) and myrLck-UD(TAMRA) to be embedded in large (LUVs) and giant unilamellar vesicles (GUVs). Instantaneous docking of protein domains in the lipid bilayer were confirmed with flotation experiments (Fig. S1). Both systems differ in size, membranous properties, and handling. Application of LUVs made it possible to maintain surface density of CD4ct and Lck-UD in the membrane at ∼415 and ∼8000 molecules per μm^2^, where the former is the reference value for Lck (50). Figure S2 presents FAM fluorescence changes in both conditions at different [Zn^2+^]_free_ values. No differences in fluorescence at 520 nm between myrCD4ct(FAM) alone or together with myrLck-UD(TAMRA) are detected for samples with a surface density of 415 molecules/μm^2^. However, for both CD4ct(FAM) alone or together with Lck-UD(TAMRA) a decrease in the fluorescence with increasing [Zn^2+^]_free_, starting at 10^−13^ M, was observed. For a higher surface density, differences are observed between donor and donor-acceptor, and it is clearly seen that the fluorescence emission pattern in the applied free Zn^2+^ concentration range is significantly different from the density 415 molecules per μm^2^.

**Figure 2.**
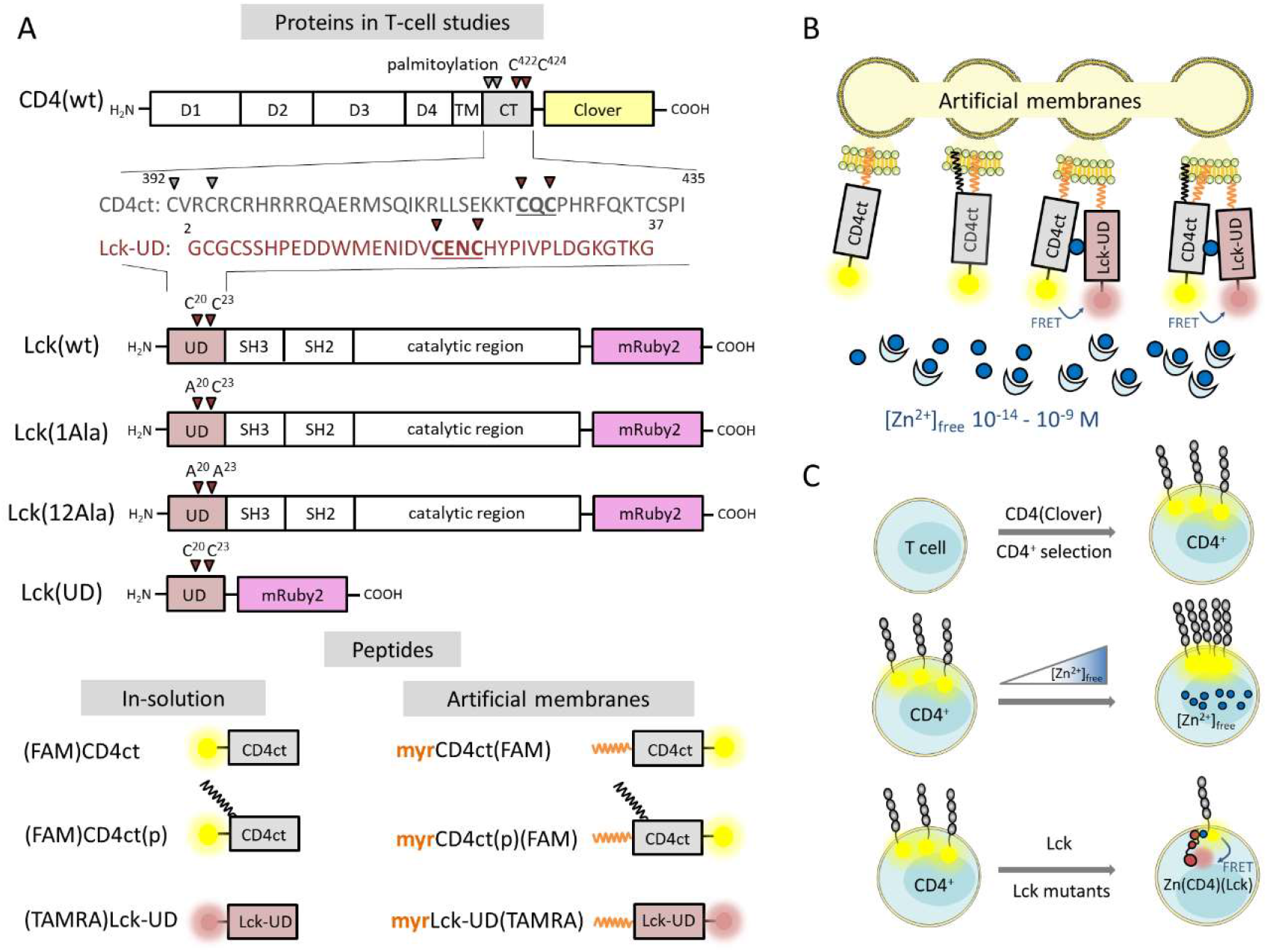
A) Schematic representation of CD4 and Lck proteins used in T-cells based studies. ct stands for cytoplasmic domain of CD4 and its amino acid sequence (CD4ct) is shown in gray. UD stands for unique domain of Lck and its amino acid sequence (Lck-UD) is shown in wine. Canonical Zn^2+^ binding cysteinyl residues are bolded and underlined. CD4 and Lck proteins were introduced as fluorescent reporters of Clover (yellow) and mRuby2 (violet) fusions, respectively. Lck(wt) mutation sites are marked as triangles and assigned. Below there is a schematic representation of model peptides used where CD4ct and Lck-UD are shown as gray and wine rectangles, respectively. FAM and TAMRA fluorescent modifications are indicated as yellow and pink blurred circles, respectively. Palmitoylation and myristylation are shown as black and orange zigzags. B) Reconstitution of CD4ct, CD4ct(p) and Lck-UD interactions in buffered free Zn^2+^ concentration (dark blue dot) in the range of 10^−14^-10^−9^ M maintained by zinc chelators (light blue half-moon) on the artificial membranes (yellow). Blurred yellow and pink circles represent FAM and TAMRA modifications, respectively. Model peptides were embedded in the membrane due to their myristylation (orange zigzag). C) Scheme of T-cell studies. CD4(Clover) was introduced to the Jurkat T-cell line and cells were selected to obtain the CD4^+^ cell line with stable overexpression of CD4(Clover). The CD4^+^ cell line was subjected to different intracellular [Zn^2+^]_free_. The CD4^+^ cell line was transiently transfected with different Lck variants to observe Zn^2+^-dependent CD4-Lck interaction studied by FRET.

**Figure 3.**
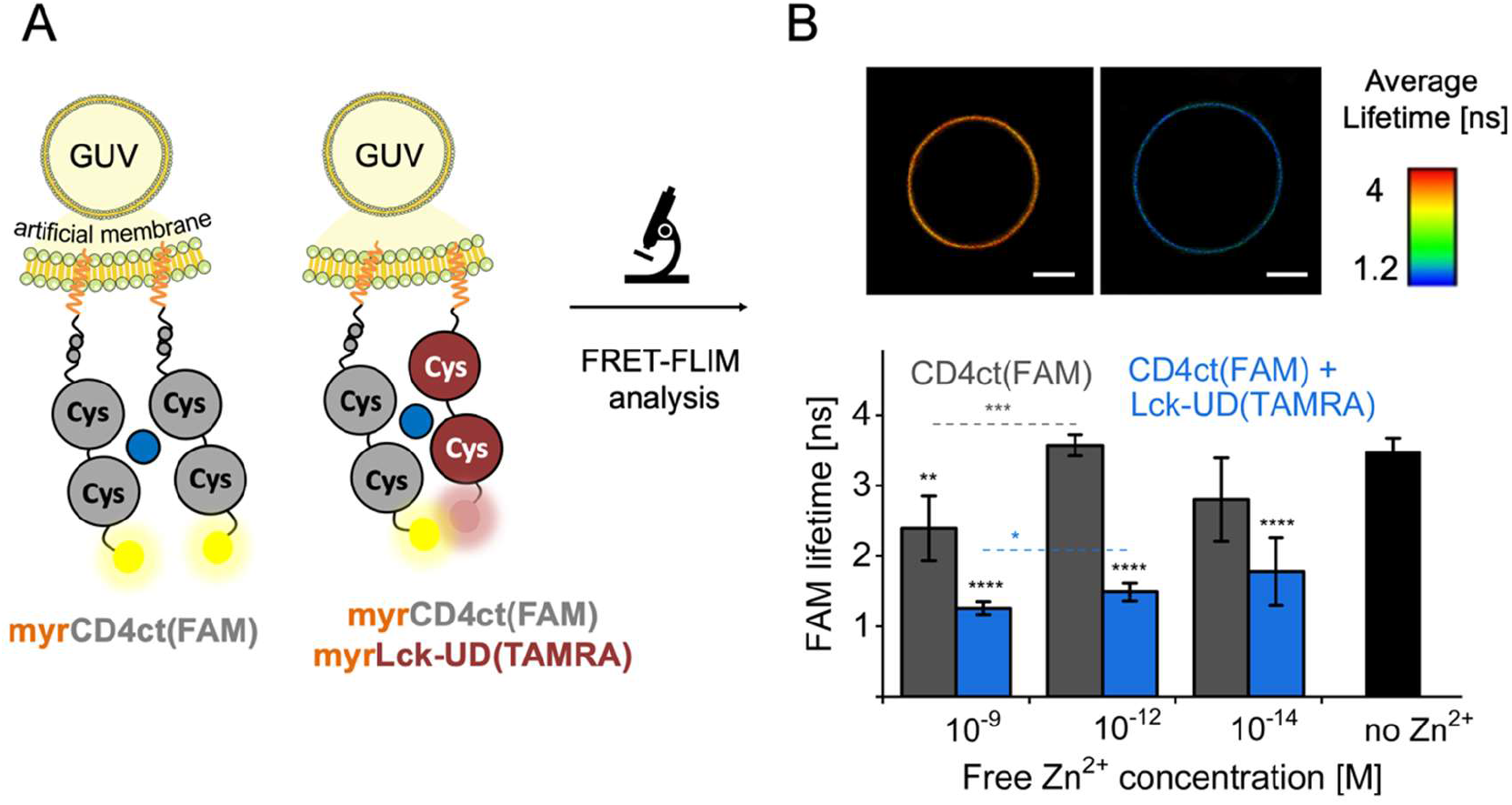
Zinc clasp assembly reconstitution on the artificial membrane. A) Liposomal membranes based on giant unilamellar vesicles (GUVs). myrCD4ct(FAM) (gray) and myrLck-UD (wine) are embedded in the membrane via myristylation at their N-terminus (orange zigzag). Cysteinyl residues within peptide domains are presented as circles. Zn^2+^ is schematically presented in blue. Fluorescent modifications at the C terminus are glowing circles. B) Representative GUV images of CD4ct(FAM) and CD4ct(FAM) with Lck-UD(TAMRA) at [Zn^2+^]_free_ of 10^−12^ M on the left and right, respectively, along with the FAM fluorescence average lifetime scale. Below there is FRET-FILM analysis of CD4ct(FAM) in gray and CD4ct(FAM) with Lck-UD(TAMRA) in blue embedded in the GUVs membrane and subjected to different [Zn^2+^]_free_. Negative control without added Zn^2+^ is shown in black. Statistical analysis was performed using one-way ANOVA with the Tukey test with p < 0.05 and n = 4-5 using Origin software.

Due to the drawbacks of LUV methodology, a GUV membrane model was here established. The major advantage is that it better resembles the cell membrane dimensions and enables one to track single liposomes under the confocal microscope. In order to observe zinc clasp formation, single GUVs that indicated the presence of both CD4ct(FAM) and Lck-UD(TAMRA) were imaged. Figure 3B shows representative GUVs of CD4ct(FAM) and CD4ct(FAM) with Lck(TAMRA) at the free Zn^2+^ concentration of 10^−12^ M, where FAM fluorescence lifetime is indicated. The results of zinc clasp reconstitution using GUVs at three different [Zn^2+^]_free_ values, 10^−9^, 10^−12^, and 10^−14^ M, show that the FAM lifetime of donor-acceptor samples differed highly significantly from control samples in all investigated free Zn^2+^ buffered conditions. Besides the observed drop of FAM fluorescence lifetime, also a difference in CD4ct(FAM) samples between free Zn^2+^ concentration of 10^−12^ and 10^−9^ M was observed. It may indicate that the Zn(CD4ct)_2_ complex is formed in addition to the zinc clasp assembly at the nanomolar free Zn^2+^ concentration.

### Palmitoylation weakens both zinc clasp assembly and CD4 dimerization

It has already been shown that the specificity of zinc clasp assembly is affected by the length of CD4ct, where the sequential and structural elements play a stabilizing role (49). Although the zinc clasp heterocomplex is prevalent when CD4ct and Lck-UD are present in equimolar amounts, the availability of additional cysteinyl and histidyl CD4ct residues turned out to play a crucial role in both complex stoichiometry and CD4ct affinity for Zn^2+^. In the cell, membrane-proximal cysteine residues undergo reversible palmitoylation; thus the influence of CD4 palmitoylation on Lck assembly was examined. Palmitoylated CD4ct(p) was compared with CD4ct in their Zn^2+^- and Lck-binding properties (Fig. 2A, 2B). Initially, relative affinities of Zn^2+^ to CD4ct, CD4ct(p) and their mixtures with Lck-UD were evaluated with the chromophoric chelating probe PAR. Upon the addition of a portion of the competitive peptide to the preformed Zn(PAR)_2_ complex, a decrease in absorbance is detected due to the Zn^2+^ transfer from the Zn(PAR)_2_ complex to the peptide(s) of choice (Fig. 4A). Both CD4ct and CD4ct(p) competition curves show a substantial decrease. The transfer of Zn^2+^ to CD4ct(p) occurs to a lower extent when compared with CD4ct, which indicates that palmitoylation may weaken the interaction of Zn^2+^ with CD4ct. A similar pattern was demonstrated in Zn(PAR)_2_ competition with CD4ct and Lck-UD equimolar mixtures where less transfer was observed to CD4ct(p) and Lck-UD (Fig. 4A). Nevertheless, it should be noted that the comparison of complexes of different stoichiometry is error prone and care should be taken in drawing conclusions. To verify the mode of Zn^2+^ binding to CD4ct and its palmitoylated version, circular dichroism was used to monitor structural changes. CD spectra of CD4ct, CD4ct(p), and their equimolar mixtures with Lck-UD indicated random coils with subtle changes at 202 and 220 nm upon Zn^2+^ addition (Fig. S3). Changes of ellipticity presented as a function of Zn^2+^-to-peptide molar ratio indicate a sharp inflection point at 1.0 for CD4ct, whereas CD4ct(p) shows a more flattened curve with the inflection point shifted towards 0.5. When CD4ct is in the equimolar mixture with Lck-UD the heterodimerization is evident, but the transition from 0.5 to 1.0 Zn^2+^-to-peptide molar ratio is noticeable in the Zn^2+^ titration curve. Palmitoylated CD4ct showed distinct behavior in the equimolar mixture with Lck-UD (ratio less than 0.5), suggesting that the assembly of a zinc clasp may not be completed in this condition.

**Figure 4.**
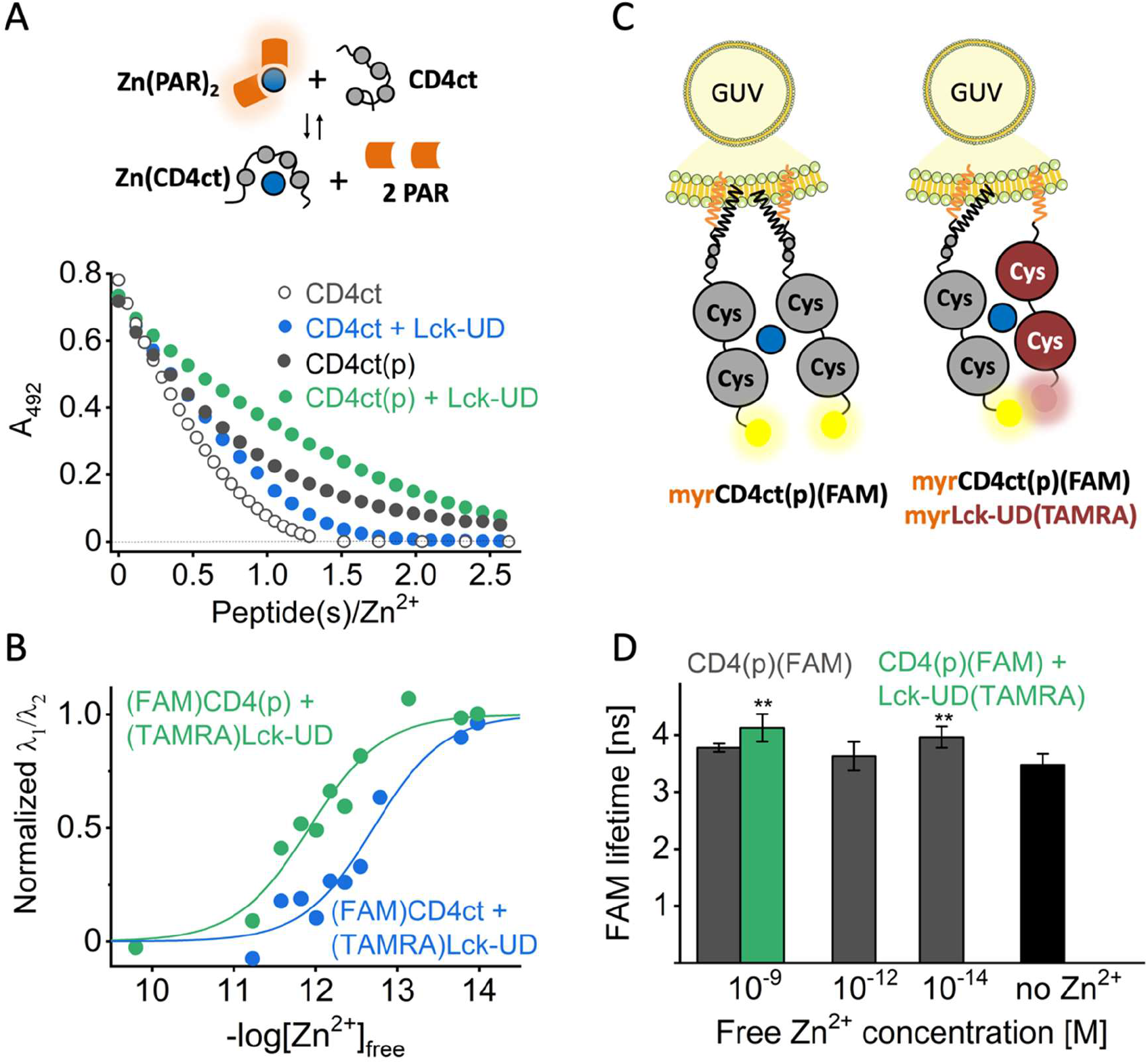
Influence of palmitoylation on zinc clasp assembly. A) Competition experiments of CD4ct (gray empty dots), CD4ct(p) (gray dots), and their equimolar mixtures with Lck-UD (blue and green, respectively) with chromophoric Zn(PAR)_2_ complex. Competition curves are presented as absorption values measured at 492 nm as a function of peptide(s) to Zn^2+^ molar ratio. B) FRET studies of fluorescently labeled CD4ct(FAM) with Lck-UD(TAMRA) and CD4ct(p)(FAM) with Lck-UD(TAMRA) are presented in blue and green, respectively. The λ_1_/ λ_2_ ratio was normalized and presented as the logarithmic function of [Zn^2+^]_free_. Data points were fitted to Hill’s function and the obtained binding isotherms are presented as green and blue lines. C) Schematic representation of a GUV vesicle with embedded CD4ct(p)(FAM) in gray and Lck-UD(TAMRA) in wine via N-terminal myristylation. Palmitoylation is depicted as a black zigzag. Zn^2+^ is represented by a blue circle. Fluorescent modification of peptide domains has been marked as yellow and wine glowing dots. D) FRET-FLIM studies of CD4ct(p)(FAM) with (green) or without Lck-UD(TAMRA) in gray embedded in the GUV membrane. Black bar indicates the control without added Zn^2+^. Lifetime of FAM fluorescence was measured for at least 10 different GUVs at different free Zn^2+^ concentrations. Statistical analysis was performed using one-way ANOVA with Tukey test with p < 0.5 and n = 3-5 using Origin software.

To evaluate differences in Zn^2+^-binding affinities, CD4ct and Lck-UD assembly was monitored in Zn^2+^-controlled media that maintain free Zn^2+^ in the range of 10^−14^ to 10^−10^ M (Table S2). Fluorescently labeled (FAM)CD4ct, (FAM)CD4ct(p) and (TAMRA)Lck-UD were mixed equimolarly and FAM (donor) and TAMRA (acceptor) fluorescence intensities were registered. Figure 4B presents Zn^2+^-binding isotherms as the function of -log[Zn^2+^]_free_ (pZn). Readouts are presented as the normalized ratio of donor to acceptor fluorescence intensity. Half-saturation pZn_0.5_ values at which half of the complex was formed were 12.7 ± and 11.9 ± 0.1 for (FAM)CD4ct and (FAM)CD4ct(p) with (TAMRA)Lck-UD, respectively. To further elucidate whether the buffered free Zn^2+^ pool influences the formation of CD4ct and Lck-UD complexes in the membrane proximity, CD4ct(p) and Lck-UD interactions were reconstituted using the GUV lipid system. Figure 4D presents the FAM fluorescence lifetime of myrCD4ct(p)(FAM) and myrCD4ct(p)(FAM) with myrLck-UD(TAMRA) that were embedded in the GUV membrane. At [Zn^2+^]_free_ of 10^−9^ and 10^−12^ M GUV liposomes with palmitoylated myrCD4ct(p)(FAM) indicated similar fluorescence lifetime as the control sample with no Zn^2+^. Contrary to myrCD4ct(FAM), no drop of FAM fluorescence lifetime was observed for myrCD4ct(p)(FAM) at the free Zn^2+^ concentration of 10^−9^ M (Fig. 3C, 4D). Moreover, when myrCD4ct(p)(FAM) was placed at the membrane surface together with myrLck-UD(TAMRA) the fluorescence lifetime did not decrease, in contrast to non-palmitoylated CD4ct.

### CD4 plasma membrane status depends on intracellular free Zn^2+^ concentration

Since the reconstitution of CD4ct, Lck-UD, and Zn^2+^ complexes on liposome membranes did show that both CD4ct and CD4ct with Lck-UD are sensitive to certain free Zn^2+^ concentrations, the CD4 presence was related to the intracellular free zinc in Jurkat cells as a model T-cell line. Intracellular free zinc concentrations were modulated by adding or depleting extracellular zinc and determined with the fluorescent zinc probe FluoZin-3AM (Fig. S4) (51). To achieve the lowest free intracellular Zn^2+^ level, the depleted media were re-supplemented with a Zn^2+^ metal buffer (SI). Altogether, the intracellular free Zn^2+^ concentrations varied from 10^−11^ to 10^−9^ M. At first, a CD4+ cell line with stable expression of CD4wt and the fluorescent reporter Clover was generated (Fig. 5a, SI). Double-positive CD4^+^ cells with different intracellular [Zn^2+^]_free_ were analyzed in terms of intra- and extracellular fluorescence (Fig. S5, Fig. 5). Figure 5B shows a histogram analysis of mean PE fluorescence (representing CD4 surface expression) in relation to the intracellular free Zn^2+^ concentration where a significant rise of the zinc signal was observed between 0.1 and 0.7 nM. Whereas fluorescence of PE was increasing with increasing [Zn^2+^]_free_, the analysis of Clover fluorescence (representing intracellular CD4) indicated different behavior. In the same range of free Zn^2+^, Clover mean fluorescence was significantly decreasing (Fig. 5C). Moreover, the changes of fluorescence occurred in a stepwise manner.

**Figure 5.**
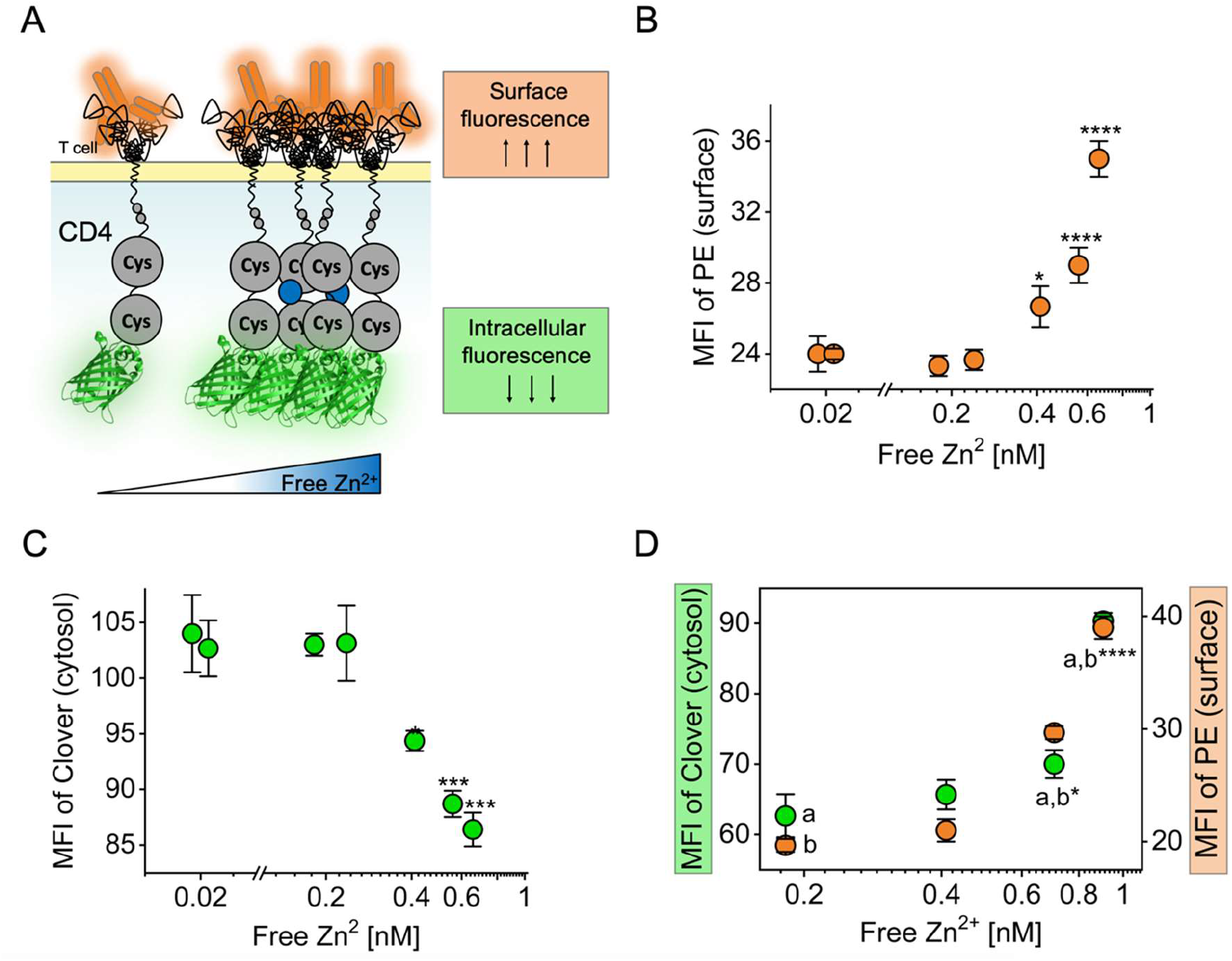
Cell surface CD4 analysis under different intracellular [Zn^2+^]_free_. A) Schematic representation of CD4+ cell line presented as a relation of anti-CD4(PE) and Clover fluorescence. B) Dot plot of Fl2-H mean fluorescence (PE) at different intracellular [Zn^2+^]_free_. C) Dot plot of Fl1-H mean fluorescence (Clover) at different intracellular [Zn^2+^]_free_. D) Intracellular staining with anti-CD4(PE). Graph presents Fl1-H (Clover) and Fl2-H (PE) mean fluorescence at different intracellular [Zn^2+^]_free_ in green and orange, respectively. Statistics were performed with one-way ANOVA with Tukey test with p < 0.5 and n = 3-5 using Origin software.

To estimate the overall level of CD4 protein in the cell with the change of [Zn^2+^]_free_, an intracellular staining protocol was introduced. Figure 5D presents the mean fluorescence of Clover and PE at different intracellular [Zn^2+^]_free_ values. It shows that the overall level of CD4 protein in the cell rises when intracellular free Zn^2+^ increases. Altogether, the observations showed that under rising intracellular [Zn^2+^]_free_ ^+^ there is a noticeable increase in the total amount of CD4 in the cell and that more of the CD4 becomes present at the cell surface.

### Assembly of CD4 and Lck depends on the cellular Zn^2+^ status

To investigate whether assembly of CD4 and Lck is dependent on Zn^2+^-binding cysteinyl residues, the CD4^+^ cell line was transiently transfected with plasmids that encode Lck with the red fluorescent reporter mRuby2 at the C terminus. Transient transfection was reported as a series of confocal microscope images 24-48 h after transfection (Fig. S6). In the study, wild-type protein Lck(wt), single and double alanine point mutants of Zn^2+^-binding cysteinyl residues Lck(1Ala) and Lck(12Ala), and the N-terminal unique domain responsible for CD4-binding via Zn^2+^ Lck(UD) were used (Fig. 2A, Fig. 6A). To monitor the CD4-Lck interaction, FACS measurements with the FRET setting were taken. CD4(Clover) and Lck(mRuby2) served as the FRET pair and the mean fluorescence of cells was recorded in three channels: donor, acceptor, and FRET. Figure 6B shows the set of FACS density dot plots where the relationship of acceptor channel fluorescence (mRuby2) to donor channel fluorescence (Clover) was depicted. Quadrant gate analysis shows that CD4^+^ cells that were transiently transfected with the Lck(wt) plasmid are mostly found in the Q2 quadrant (CD4^+,^ Lck^+^). In contrast, Jurkat cells transfected with Lck (Lck^+^) and non-transfected control (Jurkat) do not show fluorescence in the Q1 and Q2 quadrants of their dot plots.

**Figure 6.**
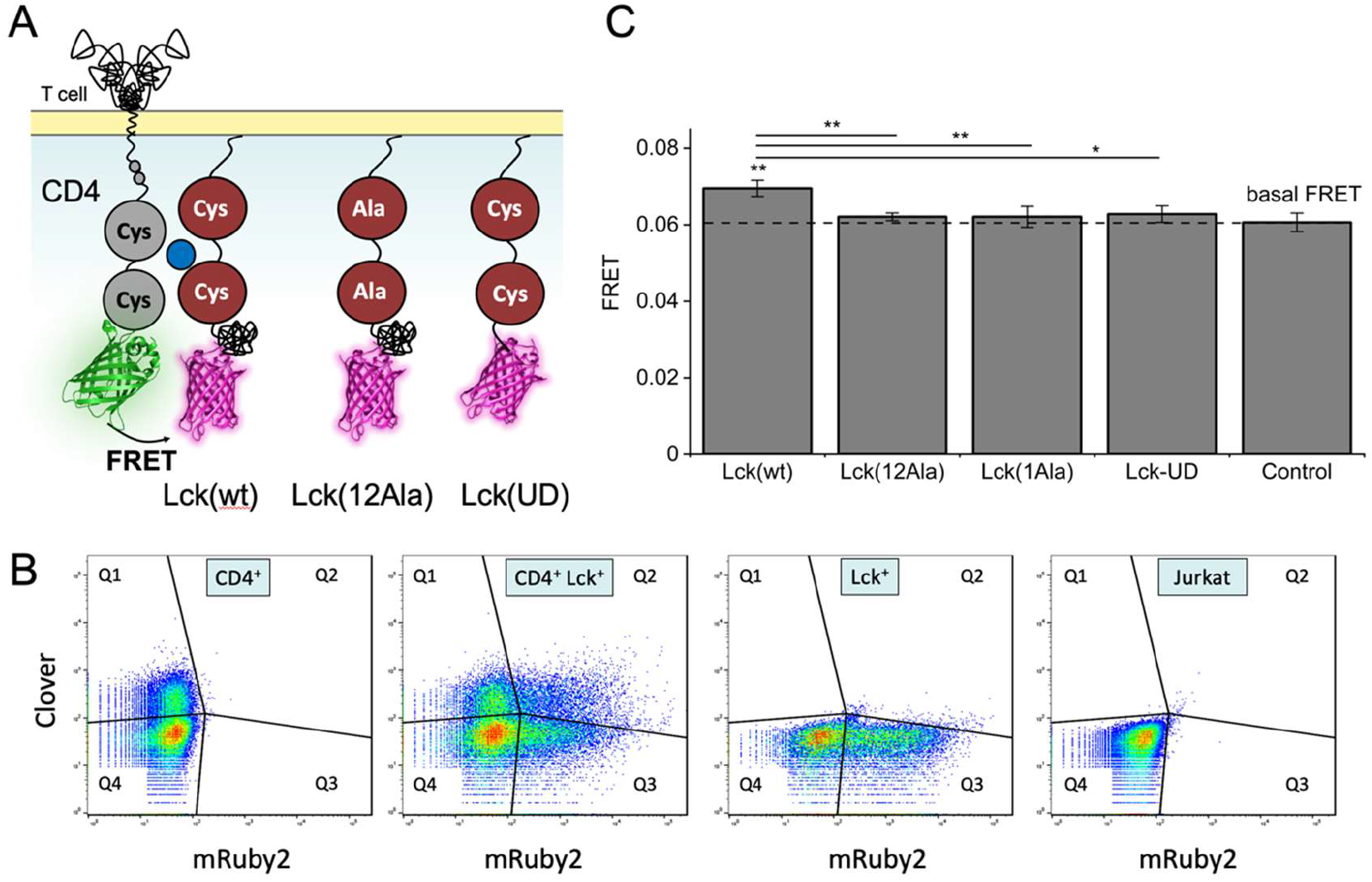
Investigation of Zn^2+^ dependence in the CD4-Lck interaction in the cell. A) Schematic representation of the zinc clasp assembly tracked by FRET between CD4(Clover) and Lck(mRuby2). CD4, Lck, and Zn^2+^ are presented in gray, wine, and blue, respectively. Zn^2+^ binding cysteinyl residues are depicted as circles. Lck(12Ala) and Lck(UD) stand for the alanine point mutation of Zn^2+^ binding cysteinyl residues and the N-terminal unique domain of Lck, respectively. B) FACS density dot plots of dual color fluorescence of Jurkat T cells. Relation of acceptor channel (mRuby2) to donor channel (Clover) is presented. The excitation of 488 and 561 nm with the emission filters of 530/30 and 610/20 nm was subjected to donor and acceptor channels, respectively. (CD4^+^) Jurkat T cells with stable CD4(Clover) expression; (CD4^+^Lck^+^) CD4^+^ Jurkat T cells transiently transfected with plasmid encoding Lck(mRuby2); (Lck^+^) Jurkat T cells transiently transfected with plasmid encoding Lck(mRuby2); (Jurkat) Jurkat T cells as a control of transfection. C) FRET studies of CD4^+^ Jurkat cell line expressing CD4(Clover) that was transiently transfected with Lck|(mRuby2) plasmids. To obtain basal FRET CD4^+^ cells were transiently transfected with mRuby2 (control). FRET was calculated as the relation of FRET to acceptor channels regarding the background fluorescence of non-transfected Jurkat cells. Statistics were performed with one-way ANOVA with Tukey test with p < 0.5 and n = 3-4 using Origin software.

For the FRET analysis, double-positive cells (Q2 quadrant) were taken and the ratio of mean fluorescence in the FRET channel to the mean fluorescence in the acceptor channel was determined. Mean fluorescence values were corrected for background fluorescence using Jurkat cell samples as a control. Figure 6C presents the results of FRET analysis of CD4^+^ cells transiently transfected with Lck(wt), Lck(12Ala), Lck(1Ala), Lck(UD), and mRuby2 as a control. A significant decrease of FRET was observed for both Lck mutants, Lck(12Ala) and Lck(1Ala), in comparison to Lck(wt). Lck(UD) also indicated a significant decrease of FRET in comparison to Lck(wt). To further examine the increase in FRET signal observed for Lck(wt), T-cell receptor signaling was activated using CD3 and CD28 (Figure S7). Upon stimulation, FRET significantly increases in CD4^+^ cells transiently transfected with Lck(wt) but not in the other Lck variants compared to the control cell line. However, the decrease of FRET in Lck(12Ala) and Lck(1Ala)-transfected cells is noticeable. Altogether, the analyses indicate that the zinc clasp is formed in the CD4^+^Lck^+^ Jurkat cell line and that its formation is related to the availability of Zn^2+^-binding cysteinyl residues.

## Discussion

Due to Zn^2+^ coordination by cysteinyl residues from both CD4ct and Lck-UD, there is a possibility for the formation of complexes of composition distinct from Zn(CD4)(Lck). For example, it has been shown that minimal domains of both CD4ct416-429 (14aa) and Lck-UD15-30 (16aa) can form homodimers in the presence of Zn^2+^ in solution (52). Further studies indicated that CD4ct396-431 binds Zn^2+^ in 1:1 stoichiometry in solution whereas the C397A mutation leads to its Zn^2+^-dependent homodimerization (21). However, in both cases, the zinc clasp heterodimer was formed predominantly.

Although the zinc clasp structure was solved 15 years ago and the participation of Zn^2+^ in its formation is indisputable, it is still omitted in the considerations on formation of the CD4-Lck complex (48). However, the importance of CD4 and Lck interaction into the process of T lymphocyte development and activation on the one hand, and the impact of Zn^2+^ on the immune system on the other, has been thoroughly investigated (25, 53-55). The reason for such disregard may reside in the indirect proof of the complex’s existence, where its disruption was observed upon treatment of cells with a membrane-permeable Zn^2+^ chelator (17). One can imagine that such treatment may cause a range of cellular Zn^2+^-involved effects that in turn cause complex dissociation, which does not indicate per se that Zn^2+^ certainly links CD4 and Lck in a cell. Another reason possibly lies in the technical difficulties of identifying metal-involved protein-protein complexes; thus their transient character makes capturing them difficult (22). The zinc clasp complex differs from common structural zinc sites in proteins, where Zn^2+^ is responsible either for polypeptide chain folding into special scaffolds (e.g., zinc fingers) or for stabilization of the whole protein or part of it. The Zn^2+^ affinity in such sites is rather high or ultrahigh and reaches low picomolar or even femtomolar *K*_d_ values, making them rather permanent under cellular conditions (56). The interprotein metal binding site occurring in the zinc clasp assembly is a different class of metal binding sites where the assembly stability from an entropic point of view is less favored due to the presence of two or more protein subunits (22). Moreover, assembly formation is strictly regulated by the actual concentrations of all its constituents (21). From the cellular perspective, the gain of an additional factor to maintain a sufficient level of interfacial Zn^2+^ complexes supports their regulatory role where assembly on demand is needed. Zinc clasp composition also raises the possibility of Zn^2+^ binding by two of the same protein subunits. The selectivity of a particular Zn^2+^ complex assembly is based mostly on the affinity of Zn^2+^ binding to the proteins, which is supported by sequential and structural elements present within CD4 and Lck (49, 52, 57).

Specificity of protein-protein interactions may be further elevated when cellular membranes are involved, phenomena observed in many cellular machineries carrying out their function (58). The involvement of distinct membrane platforms that trap or exclude proteins in induction or termination of signaling events thus has been established (59, 60-62). It may further support different local protein abundances that are used to reach the specific threshold of signaling molecules (59). To check if/how the zinc clasp complex is influenced by the presence of the membrane, two frequently used lipid bilayer models have been applied to mimic the inner leaflet of the cell membrane. While the LUV model enables control of protein densities, GUVs mimic the cellular environment better regarding size, flexibility, and stiffness of the membrane (63-65). The obtained results show that at the previously reported Lck surface density there is no specificity in the formation of the zinc clasp assembly in all investigated buffered free Zn^2+^ concentrations (50). It supports the notion that CD4ct and Lck-UD domains need spatiotemporal orchestration, and the LUV model is not sufficient. Considering that the model is limited by the interferences from unbound protein domains, the GUV system was applied, where individual liposomes were chosen to examine the interaction. Changes in the fluorescence lifetime demonstrated successful zinc clasp complex reconstitution in all applied [Zn^2+^]_free_ values from 10^−14^ to 10^−9^ M. Nevertheless, both models with embedded CD4ct indicated the decrease of fluorescence (or fluorescence lifetime) with an increase of [Zn^2+^]_free_, which may relate to the Zn^2+^-dependent CD4 dimerization.

Apart from the factor of subunit concentration and the presence of the membrane, the research undertook to verify the influence of CD4 palmitoylation on zinc clasp assembly. So far, differential protein segregation resulting from their palmitoylation has been shown to contribute to the T-cell activation processes at the cellular membrane (38, 39, 66). The zinc clasp was demonstrated to partition into lipid rafts, whereas CD4 was localized mostly in tetraspanin-enriched microdomains, and the process was found to be dependent on CD4 palmitoylation and CD4-Lck association (42, 67, 68). Conducted experiments revealed that incorporation of a long fatty acid chain on the CD4 cytoplasmic tail changes its Zn^2+^ binding properties significantly. In solution, palmitoylated CD4ct forms one-order magnitude weaker complexes with Zn^2+^ and Lck-UD. A feasible explanation is that in solution steric hindrance is substantially elevated, which prevents formation of a zinc clasp complex of higher stability. However, the interaction is completely abolished on the model membrane. Abrogated complex formation in the artificial membrane model may result from changed mobility of palmitoylated CD4. It is possible that long fatty acid chains are incorporated between membrane lipidic building blocks, making the protein domain less flexible. Such an observation also cannot exclude the scenario in which interactions in the case of CD4 palmitoylation do not appear in the investigated free Zn^2+^ concentration range. It is worth noting that the interplay of Zn^2+^ and protein concentration responsible for dimeric complex formation is also affected by their spatiotemporal orchestration within specific membrane microdomains in the cell. However, consideration of Zn^2+^ complex formation in the context of directional enrichment within membrane subdomains is still very hard. Mechanistic details of such phenomena and the potential contribution of local Zn^2+^ concentration are not yet resolved. In the face of a multitude of factors that contribute to the net outcome in the environment that would be beneficial or detrimental for complex formation, substantial efforts have been undertaken to reconstitute T-cell membrane-proximal signaling events that resulted in its most critical mechanistic details (50, 65, 69, 70). However, Zn^2+^ has never contributed to the whole picture. It seems to be high time to include it in consideration of the early events of a T-cell activation pathway.

To further explore the influence of Zn^2+^ on the CD4 membrane status its fluorescent conjugate was expressed in a Jurkat cell line. Intracellular [Zn^2+^]_free_ was modulated and proved to be in the 0.02−0.8 nM range, consistently with the previously reported levels (71, 72). Dual labeling of CD4 made it possible to observe the increase and the decrease of fluorescence in its extra- and intracellular parts, respectively, under increasing intracellular Zn^2+^ concentrations. The increase of extracellular fluorescence may be thus explained by more CD4 being present at the cellular membrane whereas the intracellular decrease may be an indication of a Zn(CD4)_2_ complex formation. However, it cannot be excluded that elevated intracellular [Zn^2+^]_free_ affected CD4 indirectly, causing its enrichment in the plasma membrane and the decrease of fluorescence due to e.g., molecular crowding. Certainly, Zn^2+^ affects the cellular behavior of CD4, an observation that has not been reported before. To support this notion, the result of intracellular staining clearly shows that the overall pool of CD4 protein rises under Zn^2+^ supplementation. Not only involvement of Zn^2+^ in the process of CD4 presentation but also its presentation itself have not yet been investigated. So far, the opposite phenomenon, i.e. CD4 internalization, has been widely studied e.g. during HIV viral infection, which positioned CD4 count as a diagnostic biomarker (73-75). Nevertheless, the presence of CD4 dimers has been shown, highlighting both its extracellular part and cytoplasmic cysteinyl residues responsible for the interaction (67, 76). The role of CD4 dimers has been proposed to be a prolonged contact with APC and increased TCR-APC avidity at the immunological synapse (76). As a result, the CD4 monomer to dimer transition may play a role in the initial engagement during T-cell activation, tuning the activation threshold.

The interplay of Zn^2+^ between CD4 and Lck is supported by the fact that the same cysteinyl residues govern CD4 dimeric assembly and the CD4-Lck formation (67). To examine zinc clasp presence in T cells, a range of Lck plasmids were introduced to the CD4+ cell line where FRET analysis showed that CD4-Lck interaction indeed occurs in the resting state of T cells (Fig. 6). Therefore, zinc status may affect the reactivity or activation threshold of resting T cells, which is seen in the reduced T-cell activity of zinc-deficient patients (77). Furthermore, this may explain why zinc supplementation has different effects on resting and activated T cells (78). The involvement of Zn^2+^ is manifested by a significant decrease of the level of the interaction when Zn^2+^ binding cysteinyl residues are mutated in Lck(wt). Interestingly, the Lck(UD) domain on its own is not able to perceive the same efficiency of the assembly as Lck(wt). It suggests that other Lck domain(s) are possibly involved and play a supporting role. Stimulation of CD4^+^Lck^+^ cells by the time matching initial phase of T-cell activation where zinc influx was found to operate (23) showed a similar effect of CD4-Lck increase. It is worth noting that CD4, Lck, and TCR molecules were reported to be present in separate clusters when cells were nonactivated, whereas upon activation larger structures were observed (79). However, large interfacial surfaces may explain the pre-activation possibility of zinc clasp formation. Therefore, Zn^2+^ may be essential for T-cell activation, but it also explains why high Zn^2+^ concentrations inhibit T-cell activation (77).

Altogether, the conducted experiments indicate that interplay between Zn(CD4)_2_ and Zn(CD4)(Lck) is feasible. However, the distinction between formation of two complexes when both are based on the same components should be made with care. On the one hand, methodological restrictions occur regarding contribution of fluorescence from two differently composed complexes. On the other, accurate determination of intracellular [Zn^2+^]_free_ is highly needed. In vitro studies demonstrate that the formation of homo- and heterodimeric Zn^2+^ complexes is well separated within the [Zn^2+^]_free_ range (49); however, the near-membrane intracellular [Zn^2+^]_free_ is not yet determined. In the view of FluoZin-3, which is generally known as a cytosolic zinc probe, it is worth noting that there are reports that it localizes within vesicles (25). But it was also found that its distribution is strictly dependent on the cell line, yet different localization has been reported (80). However, in the process of T-cell activation it has been shown that an increase in intracellular [Zn^2+^]_free_ results primarily from lysosomal Zn^2+^ release under upregulation of the ZIP8 transporter (24). Regarding the investigation of overall intracellular [Zn^2+^]_free_ it should be applicable (23), but one should keep in mind that local [Zn^2+^]_free_ values may differ.

In order to probe whether or not [Zn^2+^]_free_ changes are able to alter the molar ratio between Zn(CD4)(Lck) and Zn(CD4)_2_ species, equilibria simulations were performed based on the stability constants reported for peptide models (Fig. 7, SI) (21, 49). Apart from the fact that the stability data for Zn^2+^ complexes of peptide models could be translated quantitatively into natural protein molecules and unknown protein concentrations, the simulations show that changes in the [Zn^2+^]_free_ have a substantial impact on CD4 speciation. Taking into account CD4 and Lck at 200 and 50 nM, a change of [Zn^2+^]_free_ between 0.1 and 10 nM leads to a substantial shift towards Zn(CD4)_2_ formation (Fig. 7A, Fig. S8-S10). This notion is supported by the CD4 responsiveness to intracellular [Zn^2+^]_free_ between 0.1 and 0.6 nM observed for CD4^+^ Jurkat cells (Fig. 5). Although further increase of intracellular [Zn^2+^]_free_ could not be obtained, its higher values cannot be excluded, especially in a local manner (23). Interestingly, the fraction of Zn(CD4)(Lck) persists at ∼25% in physiologically relevant free Zn^2+^ concentrations (Fig. 7A). However, when we alter CD4, the fraction of homocomplex profile strongly depends on the free Zn^2+^ status where changes in molar ratio occur (Fig. 7B). While increasing both CD4 and Lck, at a constant ratio, increase of Zn(CD4)_2_ species occurs, with Zn(CD4)(Lck) being stable to these changes (Fig. 7C). Currently, the exact determination of CD4 and Lck level and therefore their Zn^2+^ complexes is impossible. However, local concentration of CD4-Lck is increased in the immunological synapse (10). It has also been reported that membrane signaling proteins of Jurkat cells follow diffusional trapping that leads to an increase in their concentration and/or their exclusion, thanks to the formation of distinct microdomains such as CD2/LAT/Lck coclusters (60). In addition, CD4 and the zinc clasp undergo different partitioning in the membrane subdomains and these processes were found to be correlated with TCR signaling (42, 67). It needs to be emphasized that these calculations are only approximations, and their purpose is to illustrate how dynamic the CD4/Lck system is, and that various scenarios are possible. Very likely there are optimal conditions that enhance the formation of the heterocomplex or promote formation of the homocomplex. Other processes, such as membrane partitioning and/or immune synapse formation, might also affect the local availability of CD4/Lck. Furthermore, it is possible that CD4 palmitoylation, being a reversible process, is highly involved. It is clear that a complete understanding of the role that CD4, Lck and Zn^2+^ play in T-cell activation needs further investigation.

**Figure 7.**
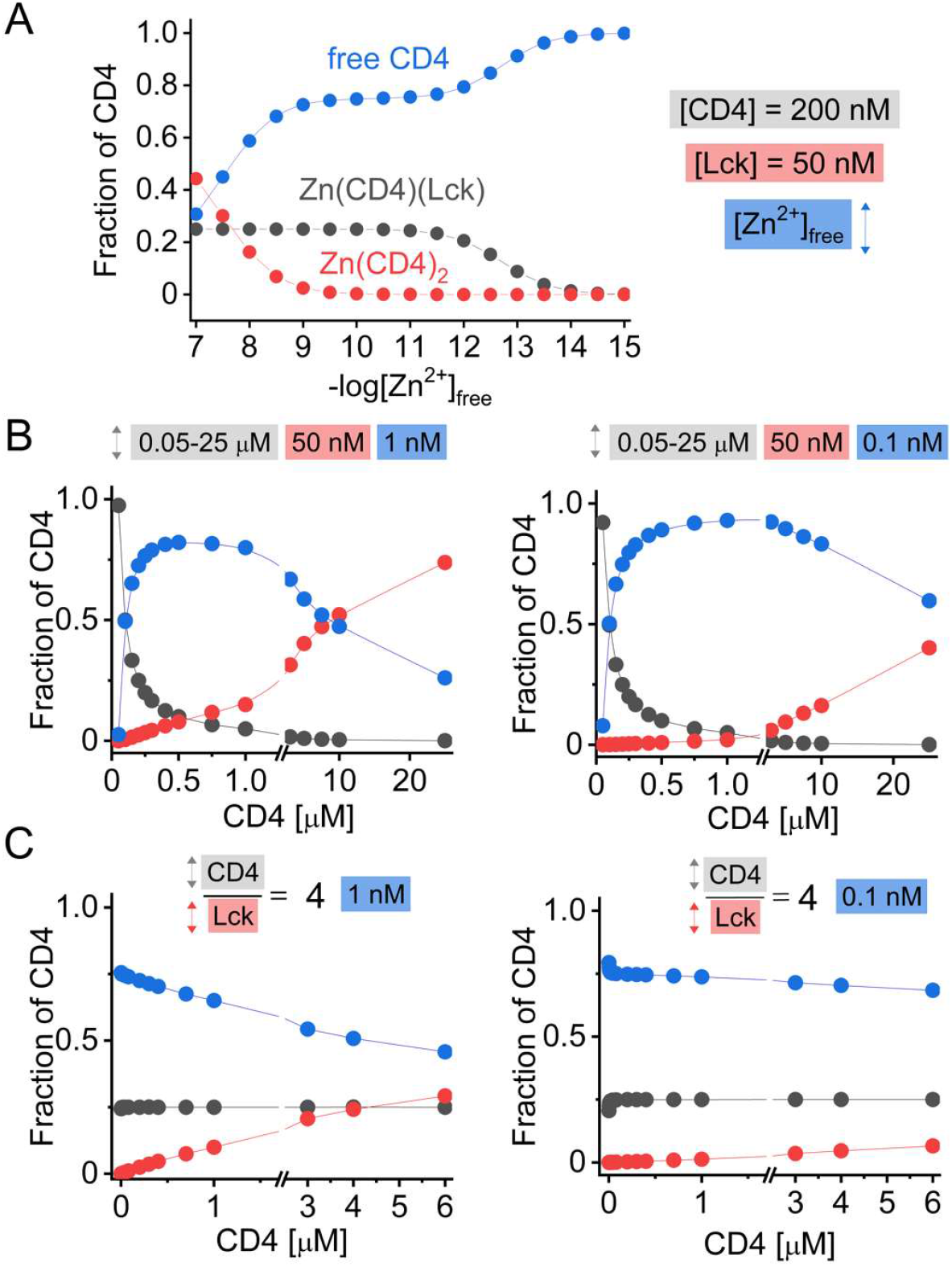
Equilibria simulation based on the stability constants obtained for Zn^2+^ complexes of peptide models presented as the fraction of CD4. Zn(CD4)(Lck), Zn(CD4)_2_ and free CD4 are presented in gray, red and blue, respectively. Concentrations of CD4, Lck, and free Zn^2+^ are shown in gray, red and blue boxes, respectively. A) Species distribution of CD4 as a function of -log[Zn^2+^]_free_. Total CD4 and Lck concentrations were set as constant values of 200 nM and 50 nM. B) Species distribution of CD4 as a function of CD4/Lck molar ratios for [Zn^2+^]_free_ at 1 and 0.1 nM. Total Lck concentration was set as 50 nM in all calculations, and CD4 varied from 50 to 25,000 nM. C) Species distribution of CD4 as a function of CD4 and Lck increasing concentrations. Total Lck concentration was increased from 0.5 to 1500 nM while CD4 increased from 2 to 6,000 nM in such a way that in all simulated points, the CD4/Lck molar ratio was set as 4.0. All distributions were calculated using HySS software (see SI).

## Materials and Methods

### Zinc clasp domain reconstitution

Zinc clasp assembly was reconstituted using LUVs and GUVs with fluorescence and fluorescence lifetime measurements, respectively. In LUV experiments, two surface densities of peptides were applied: 415 and 7412 molecules per μm^2^. In both conditions samples contained 20 nM peptides and 10 μM TCEP (50 mM HEPES, 0.1 M KNO_3_, pH 7.4) but different number of LUVs: 0.1 and 0.006 mg/ml. Peptides’ surface density was calculated with the size of one lipid 0.65 nm^2^ and 90 nm inner diameter of a liposome, which result in 52.6% of total lipids in the outer membrane surface (50, 58). Samples were incubated for 12 h prior to measurements. Concentrations of free Zn^2+^ were in the range of 10^−15^-10^−9^ M according to Table S2. Three repetitions were obtained. To visualize the zinc clasp domain in GUVs, a black 96-well glass-bottom plate (Greiner Bio-One, Kremsmünster, Austria) was blocked with 2% BSA for 30 min, and subsequently washed with 50 mM HEPES, 0.1 M KNO_3_, pH 7.4. Next, 80 μl of buffer, then peptide(s) (2 μM) and 5 μl of POPC was added to the wells, gently pipetted, and equilibrated for 15 min prior to imaging. TCEP was added at a final concentration of 5 μM. Concentrations of free Zn^2+^ were set as 10^−14^, 10^−12^ and 10^−9^ M according to Table S2.

### Competition with PAR probe

The competition of peptides with the chromophoric metal chelator PAR for Zn^2+^ (Equation 1) was performed by measuring the absorbance values of Zn(H_x_PAR)_2_ at 492 nm (ε = 71,500 M^-1^·cm^-1^) in each titration step including 3 minutes of equilibration (81). All samples contained 100 μM PAR solution partially saturated with ZnSO_4_ (10 μM) in 50 mM HEPES, 0.1 M KNO_3_, 400 μM TCEP, pH 7.4. Titration was performed in 20-30 steps up to 2.6 peptide equivalents over Zn^2+^. Stock solution of 20 mM PAR was prepared fresh in dimethyl sulfoxide (DMSO) solution.

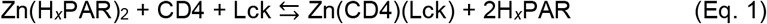

### Fluorimetry

To monitor complex formation and determine apparent dissociation constants of Zn(CD4ct)(Lck-UD) and Zn(CD4ct(p))(Lck-UD), fluorescently labeled (FAM)CD4ct, (FAM)CD4(p) and (TAMRA)Lck-UD served as FRET pairs (FAM as donor and TAMRA as acceptor). Measurements were performed at the constant temperature of 25°C with 492 nm excitation wavelength (2 nm) and 498-650 nm range in emission spectra (2 nm slit) in a quartz cuvette (Jobin Yvon Fluoromax-3 spectrofluorometer, Horiba). Peptide samples were prepared in 0.175 μM concentration (50 mM HEPES, 100 mM KNO_3_, 400 μM TCEP, pH 7.4) with zinc buffering in the [Zn^2+^]_free_ range of 10^−15^-10^−9^ M (Table S2). Samples were incubated for 12 hours and measurements were performed as three averaged scans. The ratio of donor to acceptor intensities was calculated for each experimental point and obtained data were fitted to the logarithmic version of Hill’s equation (Equation 2), where y_min_ and y_max_ refer to minimal and maximal intensities, [Zn^2+^]_0.5_ stands for free Zn^2+^ concentration where half of the complex is formed, and *n* is the cooperativity factor. Data were normalized using y_min_ and y_max_ as borders in 0 to 1 normalization.

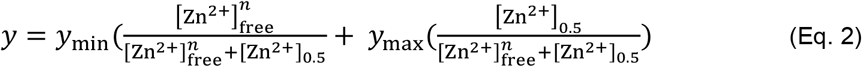

### Confocal microscopy

To determine the interaction between CD4ct and Lck-UD on the model membrane, fluorescence lifetime microscopy (FLIM), which enables quantification of FRET efficiency by measurement of the fluorescence lifetime of the donor, was used (Zeiss LSM 510 Meta). Confocal images of giant unilamellar vesicles (GUVs) that contained CD4ct(FAM) (or CD4ct(p)(FAM)) and Lck-UD(TAMRA) were gathered. For each sample, at least 10 different liposomal vesicles were chosen and imaged. Samples were excited at 488 nm for the donor and the FRET channel, and 561 nm to visualize TAMRA-containing GUVs. For donor detection, a 505 nm longpass filter was applied. Only GUVs with optimal relative fluorescence intensities of both FAM and TAMRA were imaged (40× water immersion objective with the numerical aperture of 1.2). FLIM-FRET measurements were acquired by time-correlated single-photon counting with the same microscope equipped with FLIM optics (PicoQuant). Samples were excited at 470 nm and the emission was collected with a 503-521 nm filter set. Laser power was adjusted not to exceed 900 photons/s. Images were recorded in a 512×512 format and the fluorescence lifetime was calculated for each pixel. Data analysis was done with the SymPhoTime software (PicoQuant). Each picture represented a GUV that was colored according to the average fluorescence lifetime. Data analysis of FLIM-FRET included fitting of the fluorescence lifetime decay and the instrument response function to the single exponential decay model.

### Fluorescence microscop

of CD4+ Jurkat cells was performed using the Axio Observer.Z1 with Plan-Apochromat 20× / 0.8 M27 objective (ZEISS, Oberkochen, Germany). Cells were imaged in three channels: Clover-Clover (excitation: 488 nm, emission: 525 nm filter with 25 nm bandpass) mRuby2-mRuby2 (excitation: 553 nm, emission: 605 nm filter with 35 nm bandpass) Clover-mRuby2 (excitation: 488 nm, emission: 605 nm filter with 35 nm bandpass). After transfection (48 h or 72 h) of the CD4+ Jurkat T-cell line with Lck plasmids cells were centrifuged at 4°C and half of the cells were washed with phosphate buffered saline (PBS) three times. The second half was used for FRET studies. Cells were re-suspended in 100 μl and centrifuged using a cytocentrifuge (500 × *g*, 5 min).

### FACS analysis

To analyze CD4 overexpression in the Jurkat T-cell line after transfection and during the selection with G418 antibiotic the PE-CD4 antibody was used (mouse anti-human CD4-PE clone SK3, Becton Dickinson). The level of CD4 expression level per cell and the total number of cells expressing CD4 can be obtained. To stain CD4 surface molecules 1×10^6^ cells/ml were transferred to the FACS tube, centrifuged, and washed twice with cold PBS (300 × *g*, 5 min). The pellet was re-suspended in 100 μl of cold PBS with 2% FCS. PE-labelled CD4 antibody and its isotype control (mouse IgG1,k conjugated with PE, Becton Dickinson) were added (5 μl) and incubated in the dark for 30-45 min on ice to minimize internalization. Samples were washed twice with cold PBS (300 × *g*, 5 min) and resuspended in 300 μl of PBS with 2% FCS for analysis. Statistical analysis was performed at least with three samples using one-way ANOVA and the post-hoc Tukey test with Origin software. CD4(Clover) and Lck(mRuby2) served as the FRET pair and the mean fluorescence of cells was gathered in three channels: donor (488 nm excitation, 530/30 nm emission), acceptor (561 nm excitation, 610/20 nm emission), and FRET (488 nm excitation, 610/20 nm emission). Measurements were done on FACSCalibur (Becton Dickinson) and BD FACSCanto instruments. Analysis was performed with FlowJo BD and NovoCyte Agilent software.

### Determination of intracellular free Zn^2+^ concentration

Intracellular [Zn^2+^]_free_ was measured with the specific fluorescent zinc sensor FluoZin-3. Its ester form (FluoZin3-AM) is able to permeate cell membranes, being afterwards hydrolyzed by intracellular esterases that restore FluoZin-3 that is trapped inside the cell, therefore being able to detect intracellular [Zn^2+^]_free_. The reported value of the conditional dissociation constant (*K*_d_) of the Zn^2+^-selective sensor FluoZin-3 is 8.04 nM at pH 7.4 with excitation and emission maxima at 494 nm and 517 nm, respectively (51). To determine [Zn^2+^]_free_ two borderline samples were measured, one containing 50 μM zinc chelator TPEN (*N,N,N*′,*N*′-tetrakis(2-pyridinylmethyl)-1,2-ethanediamine) and the other containing 100 μM ZnSO_4_ with 5 μM ionophore pyrithione (ZnPyr) for determination of minimal and maximal Zn^2+^ levels, respectively. The [Zn^2+^]_free_ was calculated according to the formula: [Zn^2+^]_free_ = *K*_d_ × ((F-F_min_)/(F_max_-F)) where F_min_ and F_max_ are fluorescence emission values for the samples treated with TPEN and ZnPyr, respectively. Cells were prepared in 1 ml of PBS (1×10^6^ cells). FluoZin3-AM stock solution (1 mM in DMSO) was added at a final concentration of 1 μM. Cells were incubated at 37°C for 30 min with shaking. Then cells were centrifuged (300 × *g*, 5 min), supernatant was discarded, and cell pellets were resuspended in 1 ml of PBS. The cell suspension was divided into three equal samples. For the minimal zinc sample, 7.5 μl of 2 mM TPEN was added and for the maximal zinc sample 3 μl of 10 mM ZnSO_4_ and 3 μl of 5 mM ZnPyr were added. Cells were incubated for 30 min at 37°C. Single cell fluorescence intensities were detected by FACS analysis. [Zn^2+^]_free_ was calculated as indicated above. Estimations of intracellular Zn^2+^ levels were done on non-transfected cells due to high fluorescence emission overlay. Experiments were performed in parallel with the determination of CD4 expression on the cell surface.

### Intracellular staining of CD4

Intracellular staining was performed to estimate the overall level of CD4 protein in a cell under zinc supplementation. To do so, cells must be permeabilized and fixed. Intracellular staining was done according to the manufacturer’s instructions (Intracellular Staining kit, Thermo Fisher). 100 μl of fixation buffer was added to 1×10^6^ cells/ml, vortexed and incubated for 20-30 min. Then cells were washed with 1 ml of permeabilization buffer with 2% FCS and centrifuged (300 × *g*, 5 min). Cells were re-suspended in 100 μl of Permeabilization Buffer, followed by CD4-PE antibody addition (10 μl) and incubation in the dark for 20 min. After washing with 1 ml of Permeabilization Buffer and re-suspension in 300 μl of PBS, cells were subjected to analysis.

## Acknowledgments

The research and authors (A. Krężel and A. Kocyła) were financed by the National Science Centre of Poland under Etiuda (no. 2018/28/T/NZ1/00526 to A. Kocyła), Preludium (no. 2016/23/N/NZ1/00040 to A. Kocyła) and Opus (no. 2019/33/B/ST4/02428 to A. Krężel) grants.

## Competing Interest Statement

There is no competing interest to declare.

## Author Contributions

A.K., A.C., L.R. and A.K. designed the research; A.K., A.C. and I.W. performed the research; A.K. and A.K. contributed new reagents/analytic tools; A.K. and A.K. analyzed data; A.K. and A.K. wrote the paper; A.K., I.W., L.R. and A.K. edited the paper; A.K. supervised the study.

## Notes

### Competing Interest Statement

The authors have declared no competing interest.

