## Supporting information for "A combined biochemical and cellular approach reveals Zn^2+^-dependent hetero- and homodimeric CD4 and Lck assemblies in T cells"

### Materials and methods

**Materials.** *N*,*N*-Diisopropylethylamine (DIEA), Fmoc-protected amino acids (Fmoc-Ala-OH·H2O, Fmoc-Arg(Pbf)-OH, Fmoc-Asn(Trt)-OH, Fmoc-Asp(OtBu)-OH, Fmoc-Cys(Trt)-OH, Fmoc-Gln(Trt)-OH, Fmoc-Gln(Trt)-OH, Fmoc-Glu(OtBu)-OH, Fmoc-Gly-OH, Fmoc-His(Trt)-OH, Fmoc-Ile-OH, Fmoc-Leu-OH, Fmoc-Lys(Boc)-OH, Fmoc-Met-OH, Fmoc-Phe-OH, Fmoc-Pro-OH, Fmoc-Ser(tBu)-OH, Fmoc-Thr(tBu)-OH, Fmoc-Tyr(tBu)-OH, Fmoc-Val-OH, piperidine, *O*-(benzotriazol-1-yl)-*N*,*N*,*N′*,*N′*-tetramethyluronium hexafluorophosphate (HBTU), D-biotin, [2-[2-(Fmoc-amino)ethoxy]ethoxy]acetic acid, and DL-dithiothreitol (DTT) were purchased from Iris Biotech GmbH. Trifluoroacetic acid (TFA), 1,2-ethanedithiol (EDT), thioanisole, anisole, triisopropylsilane (TIPS), COMU (1-cyano-2-ethoxy-2-oxoethylidenaminooxy)dimethylamino-morpholino-carbenium hexafluorophosphate), Fmoc-Asp(O*t*Bu)-(Dmb)-Gly-OH, Fmoc-Cys(Mtt)-OH, Fmoc-Lys(Mtt)-OH, 5(6)-carboxytetramethylrhodamine (TAMRA), 5(6)-carboxyfluorescein (FAM), guanidine hydrochloride (Gdn·HCl), myristic acid, palmitic acid, sucrose, tris(2-carboxyethyl)phosphine hydrochloride (TCEP), ethylenediaminetetraacetic acid (EDTA), HCl (trace metal grade), 4-(2-pyridylazo)resorcinol (PAR), octylglucoside, bis(β-aminoethyl ether)-*N*,*N*,*N*',*N*'-tetraacetic acid (EGTA), *N*-carboxymethyl-*N′*-(2-hydroxyethyl)-*N*,*N′*-ethylenediglycine trisodium salt ($Na_3$-HEDTA), $ZnSO_4·7H_2O$, and bovine serum albumin (BSA) were from Merck KGaA. Octyl-beta-glucoside was from Thermo Fisher Scientific. Diethyl ether, acetic anhydride, dichloromethane (DCM) and $KNO_3$ were from Avantor Performance Materials Poland S.A. Chelex 100 resin was from Bio-Rad, 4-(2-hydroxyethyl)piperazine-1-ethanesulfonic acid sodium salt (HEPES) was from Bioshop, 5,5'-dithiobis-(2-nitrobenzoic acid) (DTNB) was from TCI Europe N.V., TentaGel R RAM and TentaGel S-$NH_2$ resins were from Rapp Polymere GmbH, dimethylformamide (DMF), chloroform and acetonitrile (MeCN) were from VWR. All of the experiments were performed in chelexed buffers and solutions. All buffers were prepared with Milli-Q water obtained with a deionizing water system (Merck KGaA). POPC (1-palmitoyl-2-oleoyl-glycero-3-phosphocholine) was from Avanti Polar Lipids.

**Peptide synthesis.** Peptides were synthesized on a solid-phase support using the Fmoc strategy on TentaGel R RAM Amide Rink resin from Rapp Polymere GmbH (0.18 mmol/g substitution) and a Liberty 1 Microwave Assisted Peptide Synthesizer (CEM) according to the previous published procedures (1). Fmoc-Asp(OtBu)-(Dmb)-Gly-OH dipeptide was used for the synthesis of Lck peptide in order to prevent aspartimide formation. N-termini of peptides were acetylated with acetate anhydride (8 equiv.) or myristylated with myristic acid (10 eq.) and *N*,*N*-diisopropylethylamine (DIEA, 10 equiv). On-resin coupling of FAM and TAMRA (4 equiv.) to the N-terminus was performed with COMU (4 equiv.) and DIEA in DMF (10 equiv). GSS linker at the N terminus was applied before FAM (Table S1).

Palmitoylation of CD4ct was performed using orthogonal coupling via Fmoc-Cys(Mtt)-OH. Fmoc-Lys(Mtt)-OH was used to introduce FAM or TAMRA at the C terminus of CD4ct and Lck. The 4-methyltrityl protecting group (Mtt) was removed with 1% TFA in DCM (dichloromethane) for 15 min (5 times). Resin cleavage was performed with a mixture of TFA/EDT/thioanisole/anisole/TIPS/H2O (86:3:5:2:2:2, v/v/v/v/v/v) over a period of 4 h. After evaporation of the solution to the minimum volume under nitrogen atmosphere, peptides were precipitated with cold diethyl ether (-20°C). Crude peptide pellets were centrifuged (4°C) and washed with cold diethyl ether (5-7 times) to get rid of the excess of scavengers. Peptide crudes were stored at -20°C and non-cleaved resins after drying under a vacuum pump at 4°C.

Peptide purification was performed on an Aeris 3.6 µm PEPTIDE XB-C18 column (Phenomenex) with a gradient of acetonitrile and 0.1% TFA using the Dionex Ultimate 3000 HPLC system (Thermo Scientific). The identity of peptides was confirmed with an API 2000 Applied Biosystems ESI-MS instrument and the identified mass values with peptides sequences are listed in Table S1. Concentration of peptide stocks in 10 mM HCl was determined using a sulfhydryl group reactant, 5,5'-dithiobis(2-nitrobenzoic acid) (DTNB, $\varepsilon = 14{,}150\ M^{-1}\ cm^{-1}$ at 412 nm), prior to each experiment (2, 3). Concentrations of peptides labeled with FAM and TAMRA fluorophores were determined using fluorophores' molar absorption coefficients: $\lambda_{ex}$ (FAM) = 492 nm, $\varepsilon_{492nm} = 79{,}000\ M^{-1}\cdot cm^{-1}$ in 0.1 M NaOH and $\lambda_{ex}$ (TAMRA) = 555 nm, $\varepsilon_{555nm} = 65{,}000\ M^{-1}\cdot cm^{-1}$ in 6 M Gdn·HCl (guanidine hydrochloride) at pH 8.0 (4).

**Molecular cloning.** Nucleotide sequences of the gene encoding CD4(wt) and Lck(wt) proteins were purchased as pUC plasmids from ATG:biosynthetics GmbH, Germany (Table S3). Designed primers were terminated with nucleotide sequences encoding restriction enzyme sites (Table S4) and CD4(wt), Lck(wt), Lck(UD) were amplified using polymerase chain reaction (PCR) with Thermal Cycler T100 (Bio-Rad). PCR products were cleaned (GeneJET PCR Purification Kit, Thermo Fisher Scientific). Double-stranded DNA fragments were amplified along with expression vectors (pcDNA-mRuby2 and pcDNA-Clover, Addgene #40260 and #40259, respectively). pcDNA-mRuby2 plasmid was mutated to change the BamHI restriction site to KpnI and avoid the change of the reading frame (Table S4). In addition, two nucleotides were inserted to change the encoding linker sequence between Lck and mRuby2 from the -VP- amino acid fragment to -GTG- amino acids. Mutagenesis was performed according to the published protocol (5). Inserts and plasmids were digested at the manufacturer's conditions (4 h, 37°C), run on agarose gel (1-2%), isolated, and purified (GeneJET Gel Extraction Kit, Thermo Fisher Scientific). Inserts were then ligated with linearized plasmids using T4 DNA ligase for 1 h at 22°C (5 U/µl, Thermo Fisher Scientific). Ligation mixtures were transformed into the chemically competent *E. coli* DH5α strain. Selected colonies were cultured in LB medium with 100 µg/ml ampicillin and DNA was isolated (GeneJET Endo-Free Plasmid Maxiprep Kit, Thermo Fisher Scientific). Introduction of nucleotide sequences was confirmed with commercial sequencing (Genomed S.A., Microsynth AG).

The plasmid encoding Lck(wt)-mRuby2 was additionally mutagenized to replace $Zn^{2+}$-binding cysteinyl(s) for alanine(s); thus the 20CENC23 sequence was mutagenized to AENC and AENA, which have been abbreviated as 'Lck(1Ala)' and 'Lck(12Ala)'. Mutagenesis was performed according to the published protocol (5).

**Jurkat cell line culture.** The Jurkat T-cell line was cultured in 1640-RPMI medium supplemented with 10% fetal calf serum (FCS), 1% L-glutamine, 1% penicillin and streptomycin (Pen/Str), 1% non-essential amino acids, 1% sodium pyruvate in incubators supplied with 5% $CO_2$, 37°C and saturated humidity. The cells were examined under the microscope twice a week. The cells were counted with Trypan Blue solution according to the manufacturer's protocol and maintained at $1 \times 10^5$ viable cells per ml by dilution with fresh media. The $CD4^+$ transfected cell line was maintained with addition of G418 selective agent at 0.5 mg/ml.

**Electroporation of Jurkat cells and their selection for CD4 expression.** The day before electroporation cells were counted and adjusted to 3-5 × $10^5$ cells/ml. On the electroporation day, cells were counted and centrifuged (300 × *g*, 5 min, 7°C). For one electroporation experiment 5-10 × $10^6$ cells were washed (10 ml) and resuspended in 300 µl of cold RPMI-1640 media without FCS and Pen/Str. Cells were put into an electroporation cuvette and stored on ice for 10-20 min prior to addition of plasmids (10-20 µg) followed by a gentle shake to mix. Electroporation was performed with 300 V and 0.95 mF for 10-11 s for plasmid transfection and 8-9 s for control cells. Gentle shaking was applied before and after electroporation. After electroporation, cells were transferred to 3.5 ml of pre-warmed RPMI-1640 medium (37°C, 20% FCS, without Pen/Str) without debris. After 16-24 h, cells were diluted twice with standard RPMI-1640 medium. Cells were put under selection with 1.0 mg/ml G418 antibiotic in case of CD4 transfection or used 48-72 h after Lck transfection.

**Stimulation of Jurkat T-cell line.** After transfection of the $CD4^+$ Jurkat T-cell line with Lck (48 h or 72 h) plasmids cells were centrifuged at 4°C and washed with cold RPMI-1640 media without FCS and Pen/Str. Half of the cells were re-suspended in 300 µl of fresh medium (No FCS, No Pen/Str) and divided into 3 tubes. 100 µl of stimulation hybridoma and 50 µl of CD28 were added to each tube and incubated for 3 minutes. Stimulation hybridoma and CD28 was kindly gifted from Prof. Luca Simeoni (Molecular and Clinical Immunology, Otto von Guericke University, Magdeburg). To stop the reaction, 1 ml of cold PBS was quickly added. Cells were centrifuged (4°C, 300 × *g*, 5 min) and resuspended in 100 µl of cold PBS and stored on ice until analyzed.

**Circular dichroism spectroscopy.** Circular dichroism spectra were recorded in the range of 200 to 270 nm in 20 mM Tris, 0.1 M NaF, 0.2% octylglucoside, pH 7.4 and 300 µM TCEP (Jasco J-1500, JASCO). Peptides were in 20 µM concentration. $ZnSO_4$ was titrated up to 2 molar equivalents over peptides.

**$Zn^{2+}$-buffered system.** The series of 1 mM metal chelators (EDTA, HEDTA and EGTA) with different amounts of $ZnSO_4$ maintained $[Zn^{2+}]_{free}$ in the range of $10^{-15}$-$10^{-9}$ M. Table S2 presents $[Zn^{2+}]_{free}$ and pZn values calculated at different ratios of zinc chelator to $Zn^{2+}$ using protonation and stability constants of the chelators (6) and HySS2009 software (7). $Zn^{2+}$-to-peptide transfer during equilibration was neglected in the calculations due to the insignificant pZn changes.

**Preparation of growth media with different free $Zn^{2+}$ concentrations.** To investigate the influence of $Zn^{2+}$ on the CD4 level at the cell surface differently, $Zn^{2+}$-composed media were prepared. To chelate $Zn^{2+}$ from the media Chelex 100 resin was used (Bio-Rad). Briefly, 2.5 g of Chelex 100 per 50 ml of medium was weighed and soaked in the medium for 1 h with gentle stirring. The first portion of medium was discarded and regeneration of the Chelex 100 resin was performed. To do that 1 M HCl was added and soaked for 1 h with rolling. The fluid was discarded and the Chelex 100 resin was incubated with 1 M NaOH for 1 h with rolling. The solution was discarded and the Chelex 100 resin was washed with water for 15 minutes. Chelex resin was resuspended in water and stored at 4°C prior to use. 50 ml of media was added to regenerated Chelex 100 resin and incubated for 1 h. After the separation of the media from the resin, $Mg^{2+}$ and $Ca^{2+}$ were supplemented by the addition of 12.5 µl of 2 M $CaCl_2$ and 100 µl of 0.2 M $MgCl_2$. pH was adjusted to 7.4 and the media were sterile filtered (0.2 µm filter). Metal-free media (chelexed) were replenished with different 0.2-8 µM concentrations of $Zn^{2+}$ and a metal-buffered system was applied to obtain higher values of pZn (lower $[Zn^{2+}]_{free}$). As the $Zn^{2+}$ metal buffer 1 mM HEDTA with 2 mM $Ca^{2+}$ and 2 mM $Mg^{2+}$ was applied with the $Zn^{2+}$ concentration in the range of 0.1-0.4 mM. For the lower value of pZn (higher $[Zn^{2+}]_{free}$) media supplementation (not treated with Chelex 100 resin) was applied with $Zn^{2+}$ in the range of 0-100 µM. Cells were harvested in the prepared media for 24 h prior to analysis.

**Formation of large unilamellar vesicles (LUVs).** Liposomes were prepared by the lipid film hydration-extrusion method. POPC (1-palmitoyl-2-oleoyl-glycero-3-phosphocholine) 25 mg/ml stock solution was prepared in chloroform in a glass vial. It was stored at -20°C, tightly sealed. To prepare LUVs, 1 mg/ml POPC mixture was evaporated under nitrogen and left for 1 h at low pressure. Thin lipid films were then hydrated with 50 mM HEPES, 0.1 M $KNO_3$, pH 7.4. Five freeze/thaw cycles were performed in liquid nitrogen/warm water (40°C). The lipid mix was extruded through the polycarbonate 100 nm filter 15 times. Quality of LUVs was determined via dynamic light scattering in a concentration of 0.1 mg/ml (Zetasizer Nano ZS, Malvern Instruments, Malvern, UK). The mean diameter, polydispersity index and zeta potential of the prepared liposomes were approved. LUVs stock solution was kept at -80°C.

**Formation of giant unilamellar vesicles (GUVs).** GUVs were prepared by electroformation from 1 mg/ml stock solution of POPC in chloroform. 6 µl of POPC solution was deposited onto the electrodes (two platinum wires), followed by evaporation of the organic solvent in a vacuum desiccator for 2 h. Sucrose-containing swelling buffer (216 mM sucrose, 10 mM HEPES, pH 7.4) was added to a Teflon chamber followed by immersion of electrodes. Swelling buffer was isosmotic (240 mOsm/kg) to the experimental buffer (50 mM HEPES, 0.1 M $KNO_3$, pH 7.4). Chambers were connected to a generator ((NDN DF1641A, NDN Instrument) and exposed to a 2 V AC electric field for 2 h (1.5 h with 10 Hz, then 2 Hz). Prepared GUVs were used immediately.

**Simulations**. All calculations of species distributions were performed using HySS software (7) using $Zn^{2+}$ stability constants determined in competition experiments in our previous report (2, 8). According to that, the conditional formation constant ($\log K_{12}$) of Zn(CD4ct)(Lck-UD) and $Zn(CD4ct)_2$ is 19.47 and 14.07, respectively. In all calculations, free $Zn^{2+}$ was kept at constant values by introducing a virtual chelating agent (L) that forms with $Zn^{2+}$ a 1:1 complex (ZnL). Its apparent formation constant varied depending on the required pZn value. For instance, distribution of Zn(CD4ct)(Lck), $Zn(CD4ct)_2$, and metal-free CD4ct species was calculated by the introduction of 1.0 mM of virtual chelating agents, whose $\log K^{ZnL}$ at pH 7.4 was equal to the required pZn value. The total $Zn^{2+}$ concentration was maintained at 0.5 mM.

**Flotation experiments.** Fluorescently labeled protein domains were incubated with LUVs and applied to the sucrose gradient, followed by ultracentrifugation. The high speed of centrifugation generates LUV disks at the top of a buffer layer where docked protein domains were present. Figure 32B presents the results of a flotation experiment, where the percentage of overall fluorescence was indicated for each collected fraction. CD4(FAM) and Lck(TAMRA) samples were measured at 526 and 587 nm, respectively. Samples with myristoylated and fluorescently labeled domains possess a significantly higher percentage of fluorescence than the controls, indicating co-location of CD4(FAM) and Lck(TAMRA) in the LUV membrane model. Remaining fluorescence was observed in fraction no. 5 due to an excess of fluorescently labeled protein domain used in the experiment.

**Table S1.** Peptides obtained with standard solid-phase peptide synthesis procedures and used within the study. Sequences with corresponding observed and theoretical molecular masses are provided.

| Name | Peptide sequence | Mass obs. [Da] | Mass calc. [Da] |
|---|---|---|---|
| CD4ct | Ac-CVRCRHRRRQAERMSQIKRLLSEKKT CQCPHRFQKTCSPI-$NH_2$ | 4940.9 | 4941.6 |
| CD4ct(p) | Ac-C(palm)VRCRHRRRQAERMSQIKRLLSEK KTCQCPHRFQKTCSPI-$NH_2$ | 5180.3 | 5180.0 |
| (FAM)CD4ct | FAM-GSSCVRCRHRRRQAERMSQIKRLLSEK KTCQCPHRFQKTCSPI-$NH_2$ | 5488.4 | 5487.6 |
| (FAM)CD4ct(p) | FAM-GSSC(palm)VRCRHRRRQAERMSQIKR LLSEKKTCQCPHRFQKTCSPI-$NH_2$ | 5726.8 | 5728.4 |
| mirCD4ct (FAM) | Mir-CVRCRHRRRQAERMSQIKRLLSEKKT CQCPH RFQKTCSPI-FAM | 5652.7 | 5653.3 |
| mirCD4ct(p) (FAM) | Mir-C(palm)VRCRHRRRQAERMSQIKRLLSEK KTCQCPHRFQKTCSPIK(FAM)G-$NH_2$ | 5891.1 | 5891.8 |
| (TAMRA)Lck-UD | TAMRA-SHPEDDWMENIDVCENCHYPIVPL DGKGT-$NH_2$ | 3725.6 | 3726.1 |
| mirLck-UD (TAMRA) | Mir-GCGCSSHPEDDWMENIDVCENCHYPIVP LDGKGTK(TAMRA)G-$NH_2$ | 4476.0 | 4476.4 |

**Table S2.** Chemical components of $Zn^{2+}$-buffered media used for the $Zn^{2+}$ competition between zinc chelators and CD4 and/or Lck. Related $-\log[Zn^{2+}]_{free}$ values are presented. Chelator concentrations in each sample were 1 mM. Experiments were performed in 50 mM HEPES with $I$ = 0.1 M (from $KNO_3$) at pH 7.4 and measurements were taken at 25°C.

| $ZnSO_4$ [mM] | chelator (1 mM) | $-\log[Zn^{2+}]_{free}$ [a] |
|---|---|---|
| 0.6 | EGTA | 9.0 |
| 0.2 | | 9.8 |
| 0.05 | | 10.0 |
| 0.9 | HEDTA | 11.2 |
| 0.8 | | 11.6 |
| 0.7 | | 11.8 |
| 0.6 | | 12.0 |
| 0.5 | | 12.2 |
| 0.4 | | 12.4 |
| 0.3 | | 12.6 |
| 0.2 | | 12.8 |
| 0.1 | | 13.1 |
| 0.4 | EDTA | 13.8 |
| 0.3 | | 14.0 |
| 0.05 | | 15.0 |

[a]Protonation and stability constants of EDTA: $\beta(HL)$ = 10.17, $\beta(H_2L)$ = 16.28, $\beta(H_3L)$ = 18.96, $\beta(H_4L)$ = 20.96, $\beta(H_5L)$ = 22.47, $\beta(ZnHL)$ = 19.44, $\beta(ZnL)$ = 16.44; HEDTA: $\beta(HL)$ = 9.81, $\beta(H_2L)$ = 15.18, $\beta(H_3L)$ = 17.78, $\beta(ZnL)$ = 14.60; EGTA: $\beta(HL)$ = 9.40, $\beta(H_2L)$ = 18.18, $\beta(ZnL)$ = 12.60 (6).

**Table S3.** Sequences of purchased oligonucleotides used as forward and reverse primers in the study

| | Forward primer | Reverse primer |
|---|---|---|
| HindIII-Lck(UD)-KpnI | GCTAAAGCTTATGGGCTGTGGCTGC | TATAGGTACCTGGGGAAGCCGGC |
| HindIII-Lck(wt)-KpnI | GCTAAAGCTTATGGGCTGTGGCTGC | TATAGGTACCAGGCTGAGGCTGGTACTG |
| BamHI-CD4(wt)-EcoRI | TAGGGATCCATGAACCGGGGAGTC | GCGCGAATTCAATGGGGCTACATGTC |
| mRuby2[a] KpnI mutation | GACGATAAGGTACCGGCATGGTGTCTAAGGGCGAAGAGCTGATC | GACACCATGCCGGTACCTTATCGTCATCGTCGTACAGATCCC |
| mRuby2[a] insertion | GATAAGGATCCGGCATGGTGTCTAAGGGCGAAGAGCTGATC | GACACCATGCCGGATCCTTATCGTCATCGTCGTACAGATCCC |
| Lck(1Ala) mutagenesis | CATCGATGTGGCTGAGAACTGCCATTATCCCATAGTCCCAC | GCAGTTCTCAGCCACATCGATGTTTTCCATCCAGTCATCTTC |
| Lck(12Ala) mutagenesis | CTGAGAACGCCCATTATCCCATAGTCCCACTGGATGGCAAG | GGGATAATGGGCGTTCTCAGCCACATCGATGTTTTCCATCC |

[a]pcDNA-mRuby2 Plasmid #40260 Addgene.

**Table S4.** Sequences of purchased nucleotide sequences encoding Lck(wt) and CD4(wt).

| pUC- Lck(wt) |
|---|
| ATGGGCTGTGGCTGCAGCTCACACCCGGAAGATGACTGGATGGAAAACATCGATGTGTGTGAGAA<br>CTGCCATTATCCCATAGTCCCACTGGATGGCAAGGGCACGCTGCTCATCCGAAATGGCTCTGAGG<br>TGCGGGACCCACTGGTTACCTACGAAGGCTCCAATCCGCCGGCTTCCCCACTGCAAGACAACCTG<br>GTTATCGCTCTGCACAGCTATGAGCCCTCTCACGACGGAGATCTGGGCTTTGAGAAGGGGGAACA<br>GCTCCGCATCCTGGAGCAGAGCGGCGAGTGGTGGAAGGCGCAGTCCCTGACCACGGGCCAGGA<br>AGGCTTCATCCCCTTCAATTTTGTGGCCAAAGCGAACAGCCTGGAGCCCGAACCCTGGTTCTTCAA<br>GAACCTGAGCCGCAAGGACGCGGAGCGGCAGCTCCTGGCGCCCGGGAACACTCACGGCTCCTT<br>CCTCATCCGGGAGAGCGAGAGCACCGCGGGATCGTTTTCACTGTCGGTCCGGGACTTCGACCAG<br>AACCAGGGAGAGGTGGTGAAACATTACAAGATCCGTAATCTGGACAACGGTGGCTTCTACATCTC<br>CCCTCGAATCACTTTTCCCGGCCTGCATGAACTGGTCCGCCATTACACCAATGCTTCAGATGGGCT<br>GTGCACACGGTTGAGCCGCCCCTGCCAGACCCAGAAGCCCCAGAAGCCGTGGTGGGAGGACGA<br>GTGGGAGGTTCCCAGGGAGACGCTGAAGCTGGTGGAGCGGCTGGGGGCTGGACAGTTCGGGGA<br>GGTGTGGATGGGGTACTACAACGGGCACACGAAGGTGGCGGTGAAGAGCCTGAAGCAGGGCAG<br>CATGTCCCCGGACGCCTTCCTGGCCGAGGCCAACCTCATGAAGCAGCTGCAACACCAGCGGCTG<br>GTTCGGCTCTACGCTGTGGTCACCCAGGAGCCCATCTACATCATCACTGAATACATGGAGAATGG<br>GAGTCTAGTGGATTTTCTCAAGACCCCTTCAGGCATCAAGTTGACCATCAACAAACTCCTGGACAT<br>GGCAGCCCAAATTGCAGAAGGCATGGCATTCATTGAAGAGCGGAATTATATTCATCGTGACCTTCG<br>GGCTGCCAACATTCTGGTGTCTGACACCCTGAGCTGCAAGATTGCAGACTTTGGCCTAGCACGCC<br>TCATTGAGGACAACGAGTACACAGCCAGGGAGGGGGCCAAGTTTCCCATTAAGTGGACAGCGCCA<br>GAAGCCATTAACTACGGGACATTCACCATCAAGTCAGATGTGTGGTCTTTTGGGATCCTGCTGACG<br>GAAATTGTCACCCACGGCCGCATCCCTTACCCAGGGATGACCAACCCGGAGGTGATTCAGAACCT<br>GGAGCGAGGCTACCGCATGGTGCGCCCTGACAACTGTCCAGAGGAGCTGTACCAACTCATGAGG<br>CTGTGCTGGAAGGAGCGCCCAGAGGACCGGCCCACCTTTGACTACCTGCGCAGTGTGCTGGAGG<br>ACTTCTTCACGGCCACAGAGGGCCAGTACCAGCCTCAGCCTTGA |
| pUC- CD4(wt) |
| ATGAACCGGGGAGTCCCTTTTAGGCACTTGCTTCTGGTGCTGCAACTGGCGCTCCTCCCAGCAGC<br>CACTCAGGGAAAGAAAGTGGTGCTGGGCAAAAAAGGGGATACAGTGGAACTGACCTGTACAGCTT<br>CCCAGAAGAAGAGCATACAATTCCACTGGAAAAACTCCAACCAGATAAAGATTCTGGGAAATCAGG<br>GCTCCTTCTTAACTAAAGGTCCATCCAAGCTGAATGATCGCGCTGACTCAAGAAGAAGCCTTTGGG<br>ACCAAGGAAACTTTCCCCTGATCATCAAGAATCTTAAGATAGAAGACTCAGATACTTACATCTGTGA<br>AGTGGAGGACCAGAAGGAGGAGGTGCAATTGCTAGTGTTCGGATTGACTGCCAACTCTGACACCC<br>ACCTGCTTCAGGGGCAGAGCCTGACCCTGACCTTGGAGAGCCCCCCTGGTAGTAGCCCCTCAGT |

GCAATGTAGGAGTCCAAGGGGTAAAAACATACAGGGGGGGAAGACCCTCTCCGTGTCTCAGCTG
GAGCTCCAGGATAGTGGCACCTGGACATGCACTGTCTTGCAGAACCAGAAGAAGGTGGAGTTCAA
AATAGACATCGTGGTGCTAGCTTTCCAGAAGGCCTCCAGCATAGTCTATAAGAAAGAGGGGGAAC
AGGTGGAGTTCTCCTTCCCACTCGCCTTTACAGTTGAAAAGCTGACGGGCAGTGGCGAGCTGTGG
TGGCAGGCGGAGAGGGCTTCCTCCTCCAAGTCTTGGATCACCTTTGACCTGAAGAACAAGGAAGT
GTCTGTAAAACGGGTTACCCAGGACCCTAAGCTCCAGATGGGCAAGAAGCTCCCGCTCCACCTCA
CCCTGCCCCAGGCCTTGCCTCAGTATGCTGGCTCTGGAAACCTCACCCTGGCCCTTGAAGCGAAA
ACAGGAAAGTTGCATCAGGAAGTGAACCTGGTGGTGATGAGAGCCACTCAGCTCCAGAAAAATTT
GACCTGTGAGGTGTGGGGACCCACCTCCCCTAAGCTGATGCTGAGTTTGAAACTGGAGAACAAGG
AGGCAAAGGTCTCGAAGCGGGAGAAGGCGGTGTGGGTGCTGAACCCTGAGGCGGGGATGTGGC
AGTGTCTGCTGAGTGACTCGGGACAGGTCCTGCTGGAATCCAACATCAAGGTTCTGCCCACATGG
TCCACCCCGGTGCAGCCAATGGCCCTGATTGTGCTGGGGGGCGTCGCCGGCCTCCTGCTTTTCA
TTGGGCTAGGCATCTTCTTCTGTGTCAGGTGCCGGCACCGAAGGCGCCAAGCAGAGCGGATGTCT
CAGATCAAGAGACTCCTCAGTGAGAAGAAGACCTGCCAGTGTCCTCACCGGTTTCAGAAGACATG
TAGCCCCATTTGA

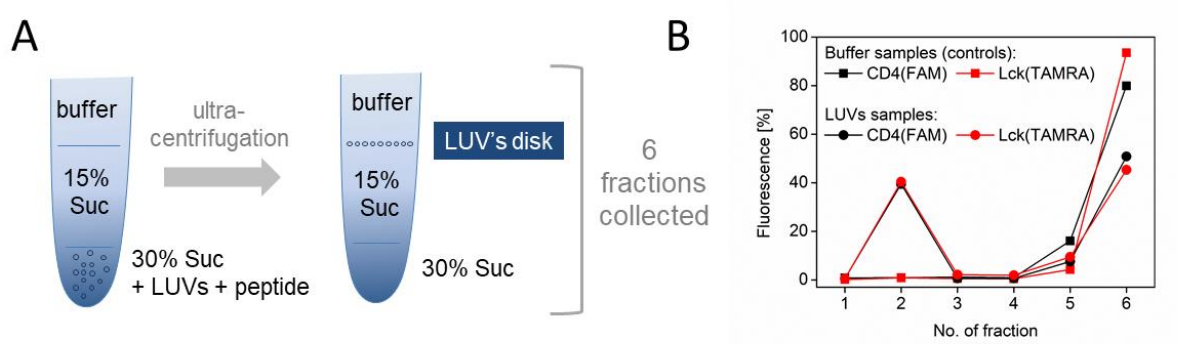


**Figure S1.** Flotation experiment of large unilamellar vesicles (LUVs) (A) LUVs with fluorescently labeled CD4(FAM), Lck(TAMRA) placed in the tube under the sucrose (Suc) gradient. Control sample consisted of buffer instead of LUVs. After ultracentrifugation on the top of the tube there is an LUV disk that freely floats together with embedded domains. Six fractions were collected counting from the top. (B) Fractions were measured fluorometrically at 526 nm (FAM) and 587 nm (TAMRA) and presented as percentage of overall fluorescence. Black and red indicate CD4(FAM) and Lck(TAMRA) samples (dots) with controls (squares).

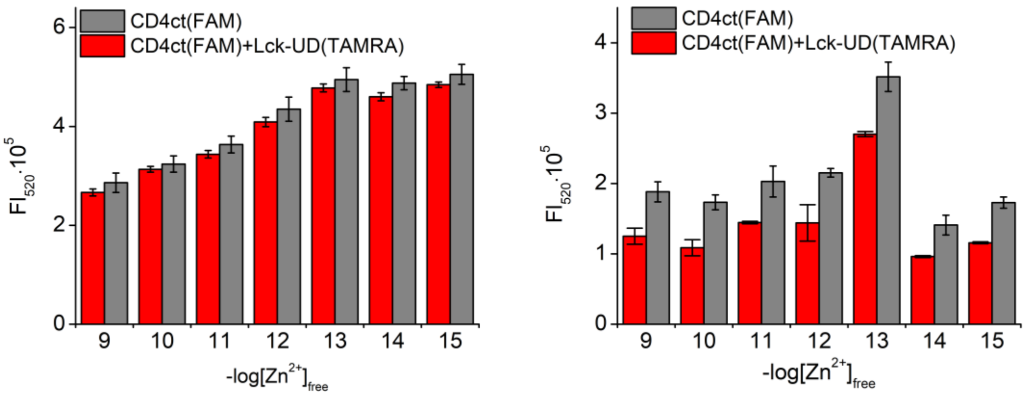


**Figure S2**. CD4(FAM) fluorescence measurements at 520 nm on the LUV model membrane with and without Lck(TAMRA) in the maintained $[Zn^{2+}]_{free}$. Two densities of embedded protein domains were studied: 415 molecules/$\mu m^2$ (left) and 7806 molecules/ $\mu m^2$ (right). In gray there are presented CD4 samples and in red the equimolar mixture of CD4 and Lck.

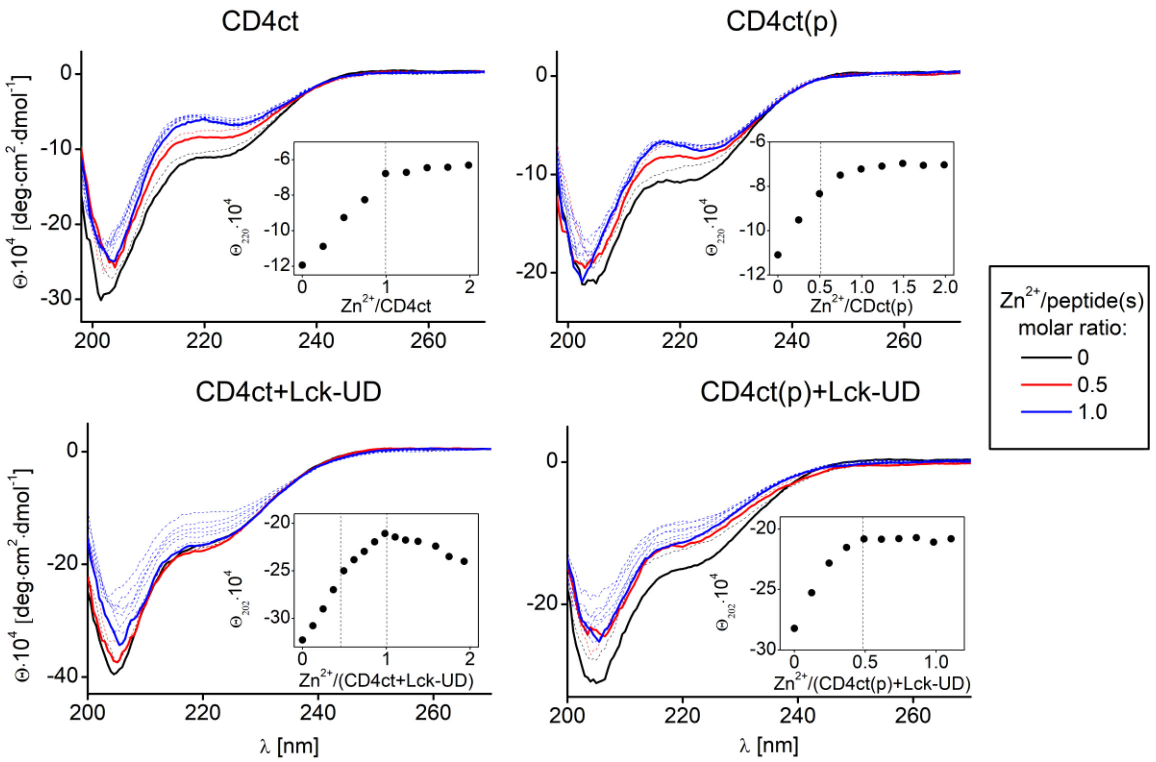


**Figure S3**. Titrations of CD4ct, CD4ct(p) and their Lck equimolar mixtures with $Zn^{2+}$ monitored by circular dichroism. CD spectra recorded in the range of 200-270 nm with the indication of 0, 0.5, and 1.0 molar ratio depicted in black, red, and blue, respectively. Changes of ellipticity at chosen wavelengths as a function of $Zn^{2+}$ to peptides molar ratio are indicated (inserts). Inflection points indicate stoichiometry of formed complexes at which structural changes are no longer observed.

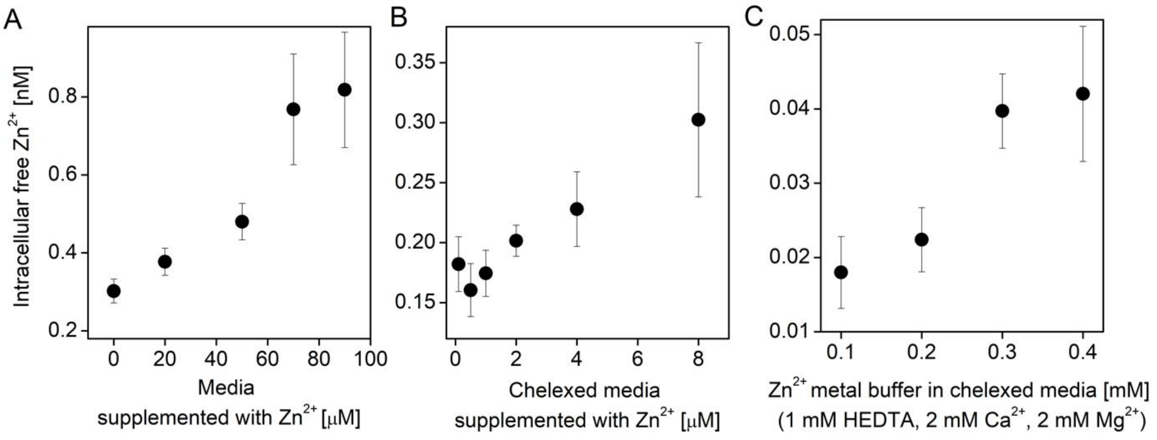


**Figure S4**. Determination of intracellular free $Zn^{2+}$ concentrations in cellular media with FluoZin-3 fluorophore. (A) $[Zn^{2+}]_{free}$ of cellular medium supplemented with $Zn^{2+}$. (B) $[Zn^{2+}]_{free}$ of chelexed cellular medium supplemented with $ZnSO_4$. (C) $[Zn^{2+}]_{free}$ of chelexed cellular medium supplemented with zinc buffer (1 mM HEDTA, 2 mM $Ca^{2+}$, 2 mM $Mg^{2+}$). Measurements were performed on three different samples.

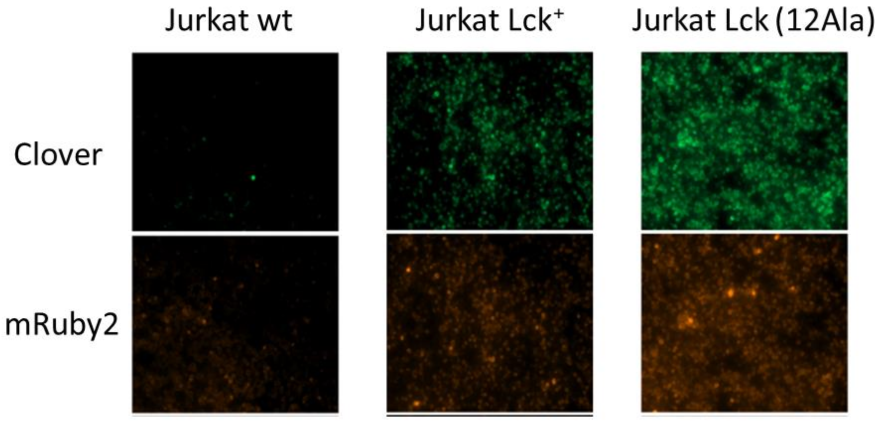


**Figure S5.** Fluorescent microscopy images of Jurkat T cells. Rows present two different channels used to observe Clover and Ruby fluorescence. Columns present three experimental outcomes: (i) non-transfected Jurkat T cells as a negative control; (ii) Jurkat T cells with CD4(Clover) expression that were electroporated with Lck(Ruby) plasmid; (iii) Jurkat T cells with CD4(Clover) expression that were electroporated with Lck(12Ala)(Ruby) plasmid.

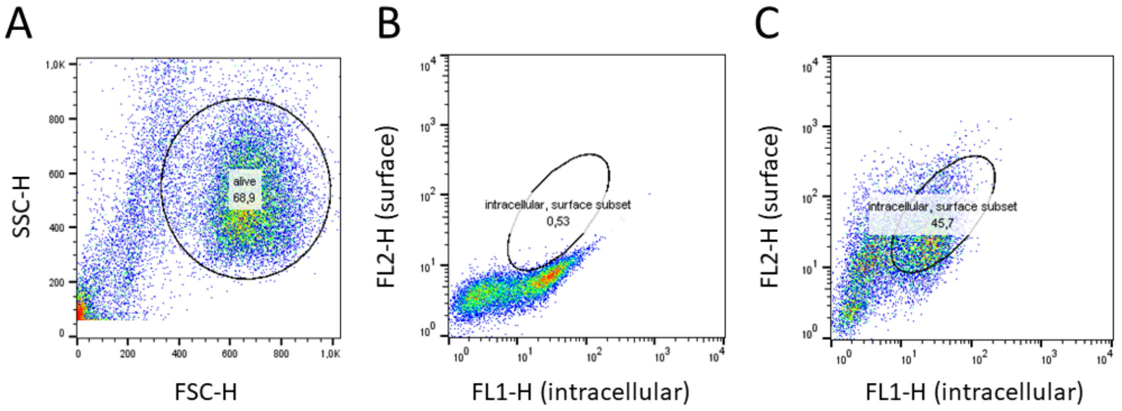


**Figure S6**. FACS cell surface CD4 analysis. (A) Dot plot of Jurkat T cells in side vs forward scatter. (B) and (C) Dot plots presented as the function of fluorescence of PE fluorophore (surface) and Clover (intracellular). Scales are shown logarithmically. (B) Isotype control staining of Jurkat T cells with CD4(Clover) overexpression. The subpopulation of Clover-expressing cells is indicated. (C) Staining of Jurkat T cells with CD4(Clover) overexpression with CD4-PE antibody. Indicated gate presents the subpopulation of double positive cells that are labeled extracellularly with PE and that express Clover protein intracellularly.

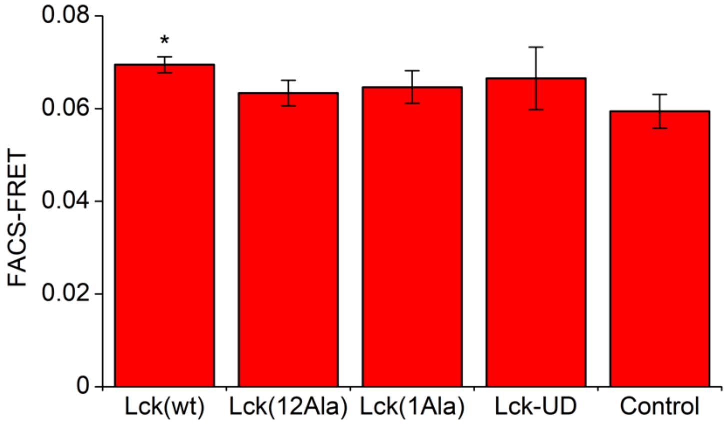


**Figure S7**. FRET studies of CD4+ Jurkat cell line expressing CD4(Clover) that was transiently transfected with Lck|(mRuby2) plasmids and subjected to 3 min of stimulation. To obtain basal FRET CD4$^+$ cells were transiently transfected with mRuby2 (control). FRET was calculated as the relation of FRET to acceptor channels regarding the background fluorescence of non-transfected Jurkat cells. Statistics were performed with one-way ANOVA with Tukey test with $p < 0.5$ and $n = 3$-4 using Origin software.

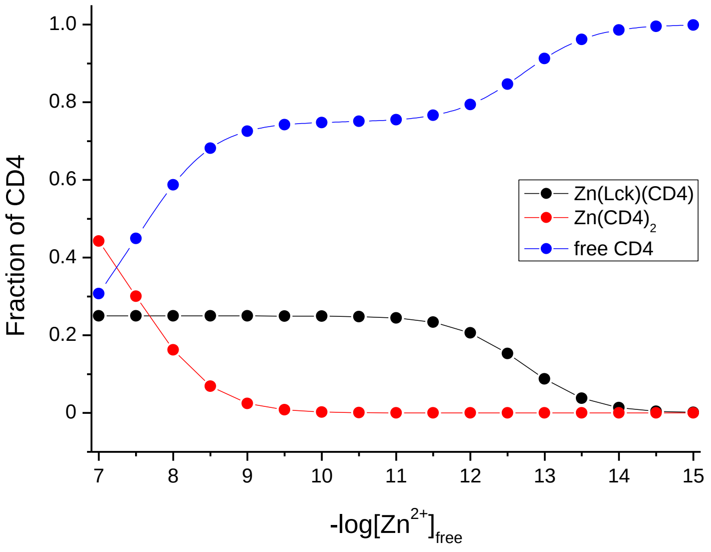


**Figure S8**. Species distribution of CD4 as a function of free $Zn^{2+}$ concentration. Total CD4 and Lck concentrations were set as constant values and were 200 nM and 50 nM, respectively. Distributions were calculated using HySS software (7).

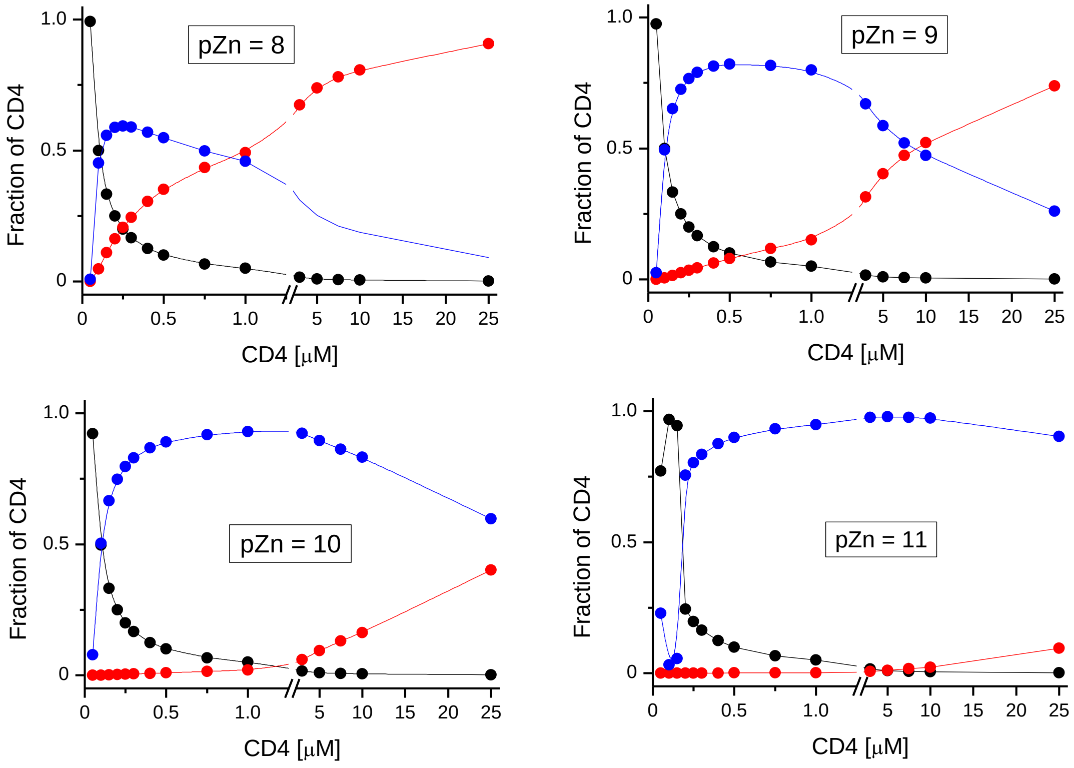


**Figure S9**. Species distribution of CD4 as a function of CD4/Lck molar ratios for four different pZn values. Total Lck concentration was set as 50 nM in all calculations, and CD4 varied from 50 to 25,000 nM. Distributions were calculated using HySS software (7).

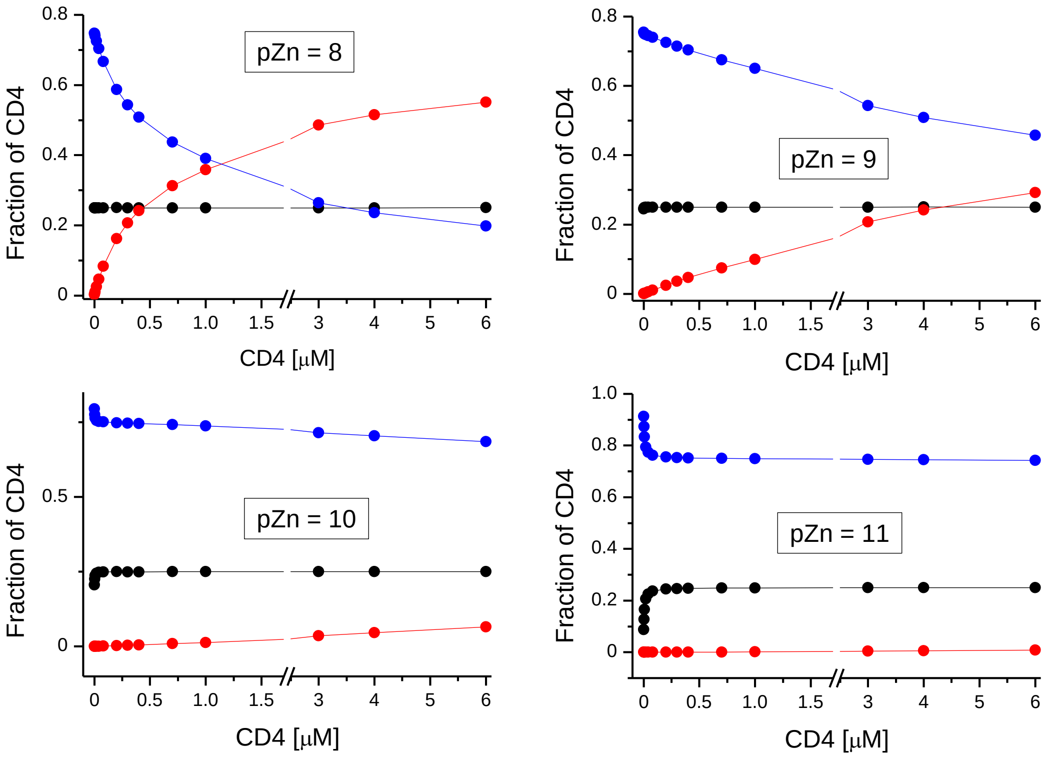


**Figure S10**. Species distribution of CD4 as a function of CD4 and Lck increasing concentrations. Total Lck concentration was increased from 0.5 to 1500 nM while CD4 increased from 2 to 6,000 nM in such a way that at all simulated points, CD4/Lck molar ratio was set as 4.0. Species distributions were calculated using HySS software (7).
